# A novel dual-chimeric surrogate virus to select, test and characterize mutants of protease and glycoprotein inhibitors

**DOI:** 10.64898/2026.01.07.698196

**Authors:** Francesco Costacurta, Andrea Dodaro, Albert Falch, Stefanie Rauch, Jakob R. Riccabona, Angelina Bäcker, Ludwig Knabl, Larry Smith, Clara T. Schoeder, Stefano Moro, Gisa Gerold, Emmanuel Heilmann

## Abstract

Antivirals are life-saving medications. However, pathogens develop resistances, mitigating their effect. Resistance studies (gain-of-function, GOF) are therefore paramount before and while antivirals are used clinically to characterize resistances and ideally study how to counteract them with other antivirals. The ability to switch from one antiviral, against which the virus is resistant, to another, depends on the availability of alternative drugs. Furthermore, targeting different parts of the virus synergistically with multiple drugs decreases the odds of simultaneous resistance. However, studying resistance, cross-resistance and remaining susceptibility requires working with the actual pathogen. Especially when introducing resistances into dangerous viruses, performing GOF studies would raise safety concerns and require firm biological containment. In this study we developed a novel dual-chimeric virus encoding both SARS-CoV-2 spike glycoprotein and main protease (M^pro^) based on the model virus, vesicular stomatitis virus (VSV). We applied selection pressure with spike and M^pro^ inhibitors, generated mutants of these viruses and determined their resistance and cross-resistance profiles by performing dose-response experiments with live and pseudotyped viruses. Additionally, we conducted *in silico* calculations to determine the impact of spike substitutions to the observed resistance phenotype. With the new technology presented here, we demonstrate that it is possible to carry out GOF studies more safely and highlight their importance in the future challenges of virological research.

## Introduction

Antiviral drugs are essential tools to combat dangerous infectious diseases. Among them, protease inhibitors have been shown to be potent as well as specific to viral enzymes in viruses such as human immunodeficiency virus (HIV-1) [1,2], hepatitis C virus (HCV) [3,4] and most recently SARS-CoV-2 [5,6]. For SARS-CoV-2, protease inhibitors proved especially effective. Two protease inhibitors, Paxlovid (nirmatrelvir/ritonavir) and Xocova (ensitrelvir), are clinically used to treat SARS-CoV-2 infections. Monoclonal antibodies (mAbs) were also brought to licensure, e.g. antibody cocktails such as EvuSheld®, and further drugs are being developed, including mAbs as well as polymerase or fusion inhibitors. Unfortunately, viruses can quickly acquire resistance when antivirals are used over longer periods, especially if they are used as single treatment rather than in combinations with inhibitors of other viral proteins [2,7–10]. This phenomenon has been documented for influenza and HIV-1 [11] and more recently, the failure of monoclonal antibodies against SARS-CoV-2 [12] and the emergence of immune escape variants with reduced susceptibility to vaccination-induced immunity. To avoid this scenario, different classes of inhibitors are often combined to engage the pathogen on multiple fronts and reduce the chance of escape mutants arising against different drugs simultaneously [13]. At the same time, resistance information is collected and stored in medical and biological databases to allow researchers and health care professionals to access highly relevant data for studying viruses or making informed treatment decisions. The data gathered in such repositories comes from clinical surveillance or prior as well as post licensing studies of a new drug. In fact, medical agencies such as the US Food and Drug Administration (FDA) require submission of clinical and non-clinical resistance studies to support a new drug application [14]. A direct approach to studying resistance-associated mutations prior to their emergence in clinical use is to investigate the authentic pathogen, such as SARS-CoV-2. The pathogen is exposed to antivirals until resistance develops. However, this kind of antiviral resistance studies are considered Gain of Function (GOF) research, which is a highly debated topic, going so far as political discussions to withdraw research funding [15,16]. Nevertheless, for pandemic preparedness and clinical treatment decision-making, resistance studies are an essential approach. To circumvent GOF research with a dangerous pathogen, we previously developed a method based on vesicular stomatitis virus (VSV). With this method, we were able to predict hallmark main protease (M^pro^) substitutions such as T21I, S144A, L167F, E166A, M49L and more, which were also described in authentic SARS-CoV-2 [17–23], among other novel substitutions not reported previously. Most importantly, we were able to predict mutations leading to these substitutions without risking accidental release of M^pro^ inhibitor-resistant SARS-CoV-2. While we focused on predicting M^pro^ mutations, other groups used VSV to predict mutations in SARS-CoV-2 spike [24–26]. The spike protein was also the first of the SARS-CoV-2 proteins to be investigated as druggable target and antigen for vaccine design. It was demonstrated that most neutralizing antibodies elicited by vaccination or infection target the receptor binding domain (RBD), which is the most immunogenic region of the spike [27,28].

The strong immunological response against the spike has inevitably put pressure on the circulating viruses, forcing them to adapt and mutate to escape the host’s defences. This, coupled with high levels of viral circulation within the population, drove SARS-CoV-2 evolution and led to the emergence of several variants, classified by the World Health Organization (WHO) according to their characteristics, such as increased transmissibility and/or enhanced immune escape [29–32]. The exposure of SARS-CoV-2 to host immune defences for prolonged periods of time – i.e. in immunocompromised individuals – can facilitate the emergence of highly mutated viruses like the Omicron variant [33]. This variant harboured 34 amino acid substitutions – 15 of which located in the RBD – rendering mAbs and immunization by vaccination or prior infection with older variants ineffective [34]. The Omicron variant outcompeted all previous variants, diverged into sub-lineages such as XBB and JN-1, prompting the update of mRNA-based vaccines [35].

Despite the pandemic being declared over in 2023, the virus continues to circulate, causing disease especially in elderly and immunocompromised individuals. Long COVID also remains as chronic condition which continues to affect people worldwide, removing them from the work force and their previous lives, silently continuing the pandemic [36]. Given this urgent clinical need and endless repertoire of new escape variants, drugs and vaccines that target more conserved regions of the spike should be developed. Additionally, pre- and post-marketing surveillance of effectiveness of drugs should always be carried out to assess and promptly gather knowledge on new viral variants in the most appropriate way.

Therefore, in this study, we expanded our safe VSV-based GOF surrogate technology to the spike protein of SARS-CoV-2. To that end, we designed a VSV variant that encodes two essential proteins of coronaviruses, which are targeted most by antiviral therapies, namely spike and M^pro^. With this dual chimeric virus, we studied mutations against licensed protease inhibitors such as nirmatrelvir and ensitrelvir or spike inhibitors such as antisera and the investigational peptide fusion inhibitor RQ-01 [37]. We showed that viruses resistant to one of the respective drugs can still be treated with a drug from a different class. With this method we could avoid dangerous GOF experiments and cross-resistance studies with authentic viruses, allowing us to investigate how viral targets might evolve under selective pressure by the deployed inhibitors.

## Results

### Design of a dual-chimeric VSV encoding SARS-CoV-2 spike and M^pro^ that is inhibited by antisera, a fusion inhibitor and protease inhibitors

To accomplish cross-resistance studies safely, we developed a novel variant of VSV that allows the selection of mutants with resistance or decreased susceptibility against different classes of inhibitors and test cross-resistance. More specifically, this virus comprises both the SARS-CoV-2 main protease (M^pro^, WA1 strain) as well as the spike protein (XBB.1.5 strain). Both transgenes were introduced into the virus such that they become essential for viral replication and propagation. This was accomplished by removing the intergenic region between the VSV glycoprotein and polymerase genes as well as the glycoprotein gene itself and replacing it with SARS-CoV-2 M^pro^ and spike glycoprotein, respectively. We generated a replicating virus and plaque purified it in Vero cells expressing two important surface proteins for SARS-CoV-2 entry, ACE2 and TMPRSS2 (Vero ↑ACE2 ↑TMPRSS2). After RNA isolation, downstream PCR amplification and sequencing of the spike protein, we detected a mutation in the furin cleavage site, a short stretch of amino acids separating the receptor binding subunit (S1) from the fusion subunit (S2) (**Supplemental Figure 1A**), that led to the substitution R682W (^681^H**R**RARS^686^ to ^681^H**W**RARS^686^). This mutation has been shown to be a cell culture adaptation event when passaging SARS-CoV-2 in Vero ↑ACE2 ↑TMPRSS2 cells [38]. To facilitate read-outs, we introduced a version of the green fluorescent protein (GFP) called mWasabi into the phosphoprotein (P) as an intramolecular insert (**Figure 1A**). The resulting virus, VSV-PmWasabi-Spike-M^pro^ (from here on referred to as VSV-S-M^pro^) can only infect susceptible cells, such as Vero ↑ACE2 ↑TMPRSS2 and HEK 293T A4 (293T ↑ACE2) cells, whereas it cannot infect standard HEK 293T cells. In contrast, the parental virus VSV-PmWasabi-M^pro^, where the native glycoprotein (G) of VSV was kept, can also infect HEK 293T as well as BHK21 cells due to the broader tropism of VSV-G (**Figure 1B**). We noticed that the dual chimeric virus replicated better at temperatures lower than 37°C, with decent replication at 34°C, which we chose for all subsequent experiments (**Figure 1C**). VSV-S-M^pro^ is a target for both fusion inhibitors and antisera as well as protease inhibitors, which inhibit spike and M^pro^, respectively. Adding either an M^pro^ inhibitor such as nirmatrelvir (NIR) or ensitrelvir (ENS), an anti-XBB.1.5 serum or the fusion inhibitor RQ-01 led to a dose-dependent inhibition of the virus, resulting in a decrease in the number of green fluorescent spots (**Figure 1D-F**). To generate VSV-S-M^pro^ mutants on 293T (↑ACE2) cells, we passaged the virus with a two-fold concentration increase of M^pro^ and spike inhibitors at each passage (NIR, ENS or RQ-01 [37]) until we reached 64- and 128-fold higher concentrations (compared to the initial dose) and then stopped incrementing it to avoid potential loss of mutated viruses.

**Figure 1.**
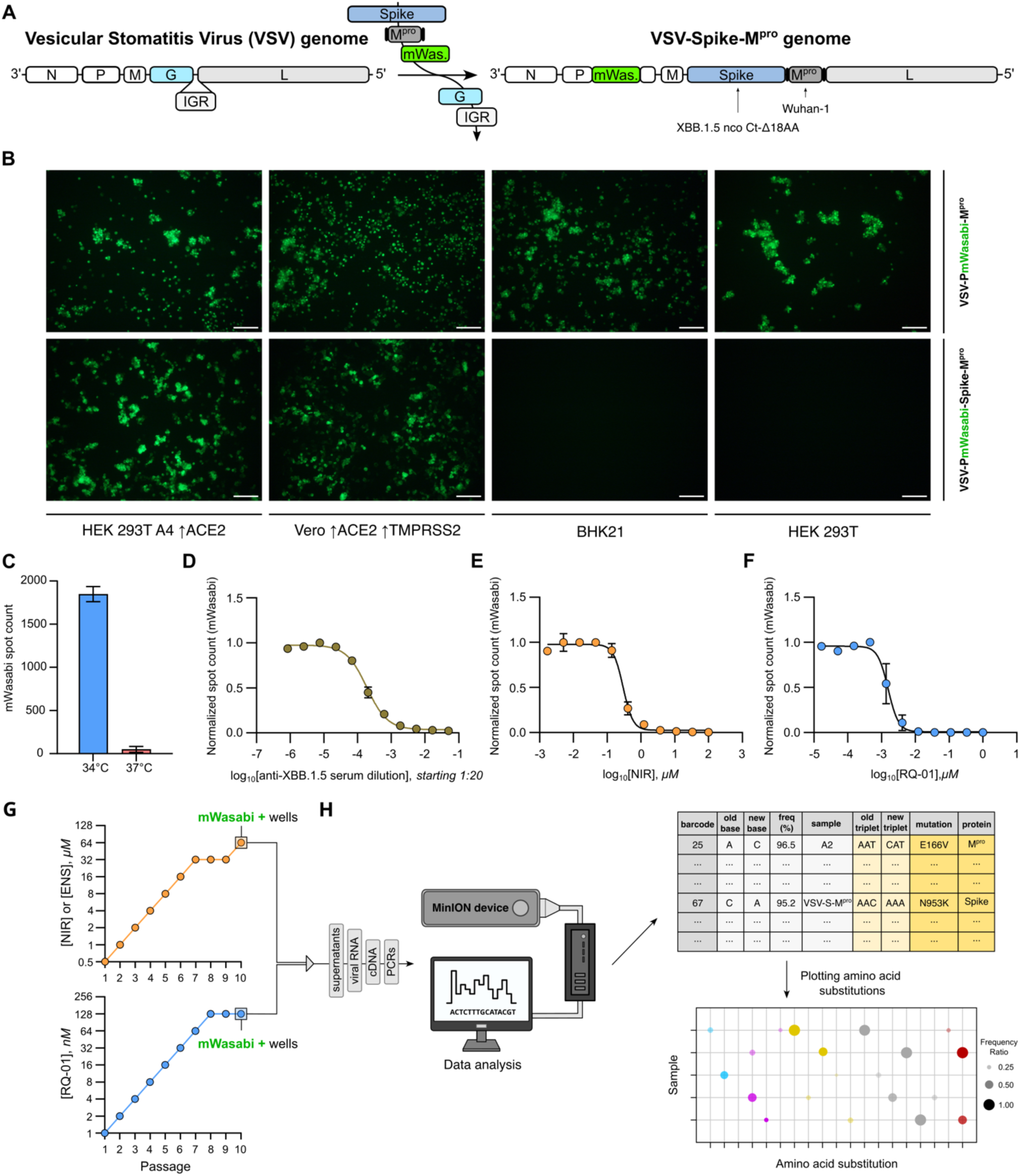
Generation and inhibition of a dual-chimeric VSV encoding SARS-CoV-2 spike and M^pro^ and selection pressure experiments workflow. **A)** Schematic representation of the genomes of VSV and VSV-S-M^pro^. We replaced VSV-G with the C-terminally truncated (18 amino acids), non-codon optimized SARS-CoV-2 XBB.1.5 spike variant. We replaced the intergenic region (IGR) between G and L with the SARS-CoV-2 M^pro^ Wuhan-1 strain. We added mWasabi as marker gene to monitor viral replication more easily as shown in panel B. **B)** Representative microscopy pictures of cells infected with either VSV-M^pro^ (here indicated as VSV-PmWasabi-M^pro^ to highlight the marker gene) or VSV-S-M^pro^ (indicated as VSV-PmWasabi-Spike-M^pro^) at two days post infection obtained at 100X magnification. Scale bar: 100 μm. Both viruses infect HEK 293T-A4 ↑ACE2 and Vero ↑ACE2 ↑TMPRSS2 cells, whereas only VSV-PmWasabi-M^pro^ infects BHK21 and HEK 293T cells as well. **C)** Representative bar plot showing the difference in mWasabi spot count measured two days post infection at either 34 or 37°C. Data are shown as mean ± SEM of four technical replicates. **D)** Representative dose-response experiments with an anti-XBB.1.5 spike serum. The highest dose started at 1:20 dilution of the serum sample and then serially diluted in a three-fold manner. mWasabi spot count was measured at 72 hours post infection. Data are presented as mean ± SEM of two technical replicates. **E)** Representative dose-response experiments with the M^pro^ inhibitor NIR. mWasabi spot count was measured at 95 hours post infection. Data are presented as mean ± SEM of two technical replicates. **F)** Representative dose-response experiments with the fusion inhibitor RQ-01. mWasabi spot count was measured at 96 hours post infection. Data are presented as mean ± SEM of four technical replicates. **G)** Top: we passaged parental VSV-S-M^pro^ on 293T-A4 ↑ACE2 cells with a two-fold concentration increase of M^pro^ inhibitors at each passage (NIR or ENS). Bottom: we passaged parental VSV-S-M^pro^ and a VSV-S-M^pro^-E166V variant with a two-fold concentration increase of RQ-01 at each passage. **H)** Workflow for downstream ONT sequencing and data analysis.

We initially selected VSV-S-M^pro^ variants of M^pro^ using inhibitors NIR and ENS. We detected a few M^pro^ mutations, among which we found E166V, a hallmark resistant substitution of M^pro^ already reported by several research groups [39–41]. We further passaged VSV-S-M^pro^ parental and a VSV-S-M^pro^-E166V variant (N-A2.2) with the fusion inhibitor RQ-01 (**Figure 1G**). In both cases, we collected supernatants, purified genomic RNA, transcribed cDNA and performed PCRs on the region of interest as well as Oxford Nanopore (ONT) sequencing to identify mutations and display the associated amino acid substitutions (**Figure 1H**).

### Selected VSV-S-M^pro^ main protease variants are resistant to NIR and ENS but not to BOF, anti XBB.1.5 serum and RQ-01

We detected several M^pro^ mutants by passaging parental VSV-S-M^pro^ with its specific inhibitors NIR (**Figure 2A**) and ENS (**Figure 2B**), as outlined above. Out of the pool of M^pro^ mutants we plaque purified variants aiming for previously described hallmark mutants at position E166 [19,23,39–41], such as E166V and E166K (**Figure 2A,B**). We tested the first panel of M^pro^ mutants against NIR and an anti-XBB.1.5 serum. As expected, passaging with M^pro^ inhibitors did not affect the antiserum’s efficacy, whereas NIR’s inhibitory activity decreased. (**Figure 2C,D**). We tested a second panel of M^pro^ mutants against NIR and the fusion inhibitor RQ-01 and found that M^pro^ mutations did not confer resistance nor cause a decrease in the susceptibility to RQ-01 (**Figure 2E,F**). We additionally tested this same panel of mutants against NIR, ENS and another M^pro^ inhibitor that had been shown to retain activity against M^pro^ resistant variants, named bofutrelvir (BOF) (**Figure 2G,H**) [42]. BOF successfully inhibited all variants.

**Figure 2.**
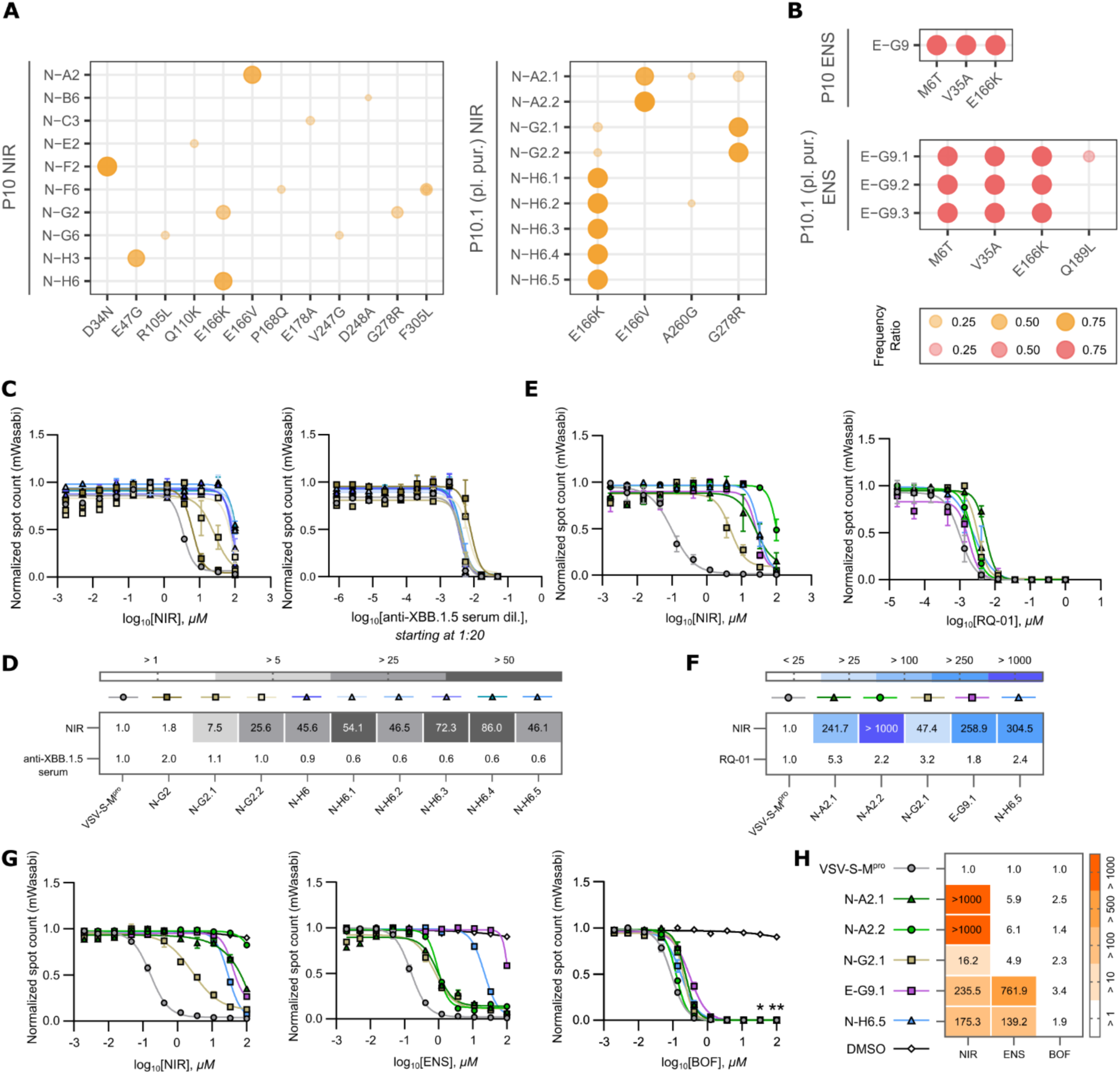
Selected VSV-S-M^pro^ variants are resistant to NIR and ENS but not to BOF, anti XBB.1.5 serum and RQ-01. Graphs in **A** and **B** provide the following information: passage (left side of the graph), sample name, amino acid (AA) substitution(s) of each sample and the population fraction/frequency ratio of the sequence carrying the single nucleotide variant (SNV) that leads to that specific AA substitution (from 0 to 1). The AA substitutions are plotted on the x-axis, and the sample names are plotted on the y-axis. Sample nomenclature: indication of the inhibitor used to select mutants (N- for NIR; E- for ENS); indication of the well from which the sample was collected (A2, H6, etc…) and for plaque-purified samples, indication of the plaque number (i.e. N-A2.1, N-A2.2, stand for two different plaques picked from the same virus sample). **A)** ONT sequencing results of samples collected after passaging 10 times with NIR (left) and after plaque purification (right). **B)** ONT sequencing results of the only sample collected after passaging 10 times with ENS (top) and after plaque purification (bottom). Frequency ratio legend: small and faded circles indicate a low population/frequency ratio of specific mutants in the sequences pool from a specific sample, big and vivid circles indicate the opposite. **C**) Dose-response curves of NIR (left) and anti-XBB.1.5 serum (right) with the first panel of M^pro^ variants (sample N-G2 and plaques G2.1 and .2; sample N-H6 and plaques H6.1, .2, .3, .4 and .5) selected during serial passaging with NIR. MOI = 0.1. mWasabi spot count was measured at 96 hours post infection. Data is presented as mean ± SEM of two technical replicates per condition. Symbols and lines of the dose-response curves are in **panel D**, above the related VSV-S-M^pro^ variant data. **D)** Heat map of the IC_50_ fold-changes of all the tested VSV-S-M^pro^ variants compared to the parental virus, for both NIR and anti-XBB.1.5 serum. **E)** Dose-response curves of another subset of M^pro^ variants (samples N-A2.1, N-A2.2, N-G2.1, E-G9.1 and N-H6.5) against the inhibitors NIR (left) and RQ-01 (right). MOI = 0.01. mWasabi spot count was measured at 83 hours post infection. Data is presented as mean ± SEM of four technical replicates per condition. Symbols and lines of the dose-response curves are in **panel F**, on top of the related VSV-S-M^pro^ variant. **F)** Heat map including the calculated IC_50_ fold-changes of all the tested VSV-S-M^pro^ variants compared to the parental virus, for both inhibitors. **G)** Dose-response experiments of the same panel of variants (**E** and **F**) against the M^pro^ inhibitors NIR (left), ENS (middle), BOF (right). MOI = 0.01. mWasabi spot count was measured at 61 hours post infection. Data is presented as mean ± SEM of three/four technical replicates per condition. Symbols and lines of the dose-response curves and M^pro^ variants are displayed in **panel H**, beside the related VSV-S-M^pro^ variant. Dose-response experiments of the same subset of VSV-S-M^pro^ variants against NIR (left), ENS (middle) and BOF (right). **H)** Heat map including the calculated IC_50_ fold-changes of all the tested VSV-S-M^pro^ variants compared to the parental virus, for all the inhibitors. * BOF shows cytotoxicity at ∼33 mM, ** BOF is highly cytotoxic at 100 mM [18].

### Selected VSV-S-M^pro^ spike HR1/HR2 variants display decreased susceptibility to RQ-01

After assessing our system’s capability to generate protease inhibitor-resistant M^pro^ variants, we sought to assess the same by serially passaging the parental VSV-S-M^pro^ and the N-A2.2 variant (M^pro^-E166V) with the fusion inhibitor RQ-01. We passaged both viruses with the aim of selecting spike-only as well as spike and M^pro^ variants, respectively. After 10 passages, we sequenced our pool of selected viruses and detected several substitutions (**Supplemental Figure 1B,C**). Because RQ-01 acts by blocking viral fusion mediated by heptad repeat 1 and 2 domains (HR1, HR2), we expected to find multiple viruses harboring substitutions within these two domains. We then decided to restrict the more detailed investigation mainly to variants with different combinations of HR1-only, HR2-only or HR1/HR2 substituted variants (**Figure 3A,B**). We observed that viruses with substitutions in both M^pro^ and spike proteins showed that the resistance against NIR could be very pronounced, whereas the susceptibility against RQ-01 was comparatively less reduced (**Figure 3C**). Viruses with spike-only substitutions generally remained susceptible to NIR. RQ-01 is a mimetic of the HR2 domain of the spike 6-helix bundle, which is essential in the structural re-arrangements during viral membrane fusion. Mapping some of the selected amino acid substitutions onto the crystal structure of the 6-helix bundle shows that substitutions occur both in the HR1 domain as well as the HR2 domain (**Figure 3D**).

**Figure 3.**
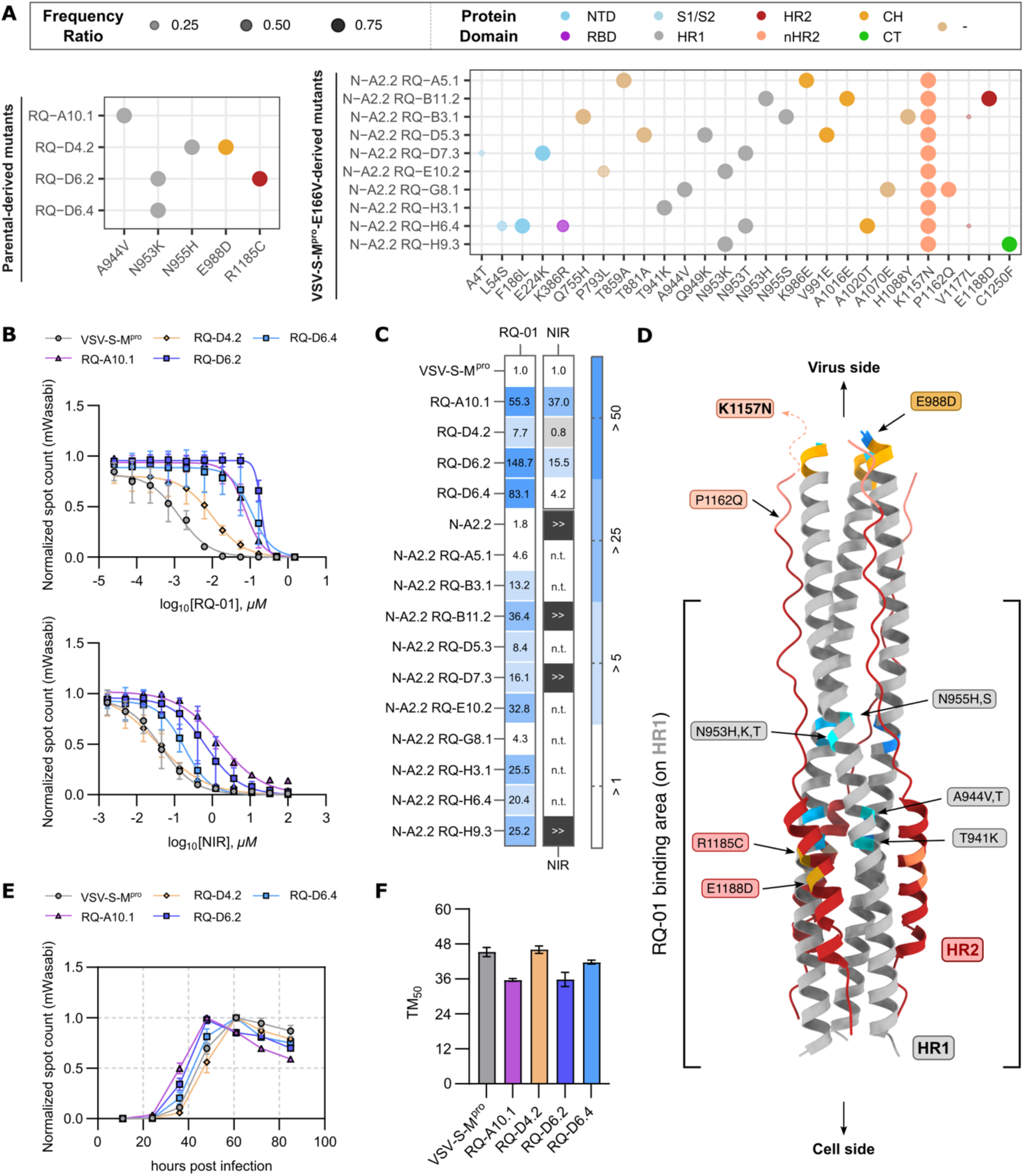
ONT sequencing, 3D structure annotation and IC_50_ fold-changes of selected VSV-S-M^pro^-derived HR1/HR2 variants after plaque purification. **A**) Top: Legend box indicating the frequency ratio. Small and faded circles indicate a low population/frequency ratio of specific mutants in the sequences pool from a specific sample, whereas big and vivid circles indicate the opposite. Color-coding is assigned according to the spike protein domains [43]. “CH” = central helix; “HR1” = heptad repeat 1; “HR2” = heptad repeat 2; “NTD” = amino-terminal domain; “RBD” = receptor binding domain; “-” = no domain assigned and S1/S2 = furin cleavage site. “nHR2” = near HR2 is annotated to highlight two substitutions in proximity to the HR2 domain (K1157N and P1162Q). Bottom: annotation of AA substitutions in the spike protein of viruses selected with RQ-01, from parental VSV-S-M^pro^ (left) and VSV-S-M^pro^ N-A2.2 (right). We did not include R682W in panel **A** as it is present in all viral sequences. **B**) Dose-response curves for the parental virus VSV-S-M^pro^ and some spike mutants RQ-A10.1, RQ-D4.2, RQ-D6.2 and RQ-D6.4 with RQ-01 (top) and NIR (bottom). Data is presented as mean ± SD of 2 independent experiments. **C)** Heat map including IC_50_ fold-changes of all spike variants vs NIR and RQ-01 compared to the originating variants VSV-S-M^pro^ and VSV-S-M^pro^-E166V (N-A2.2). “n.t.” = not tested. Data is presented as mean of 2 independent experiments, except for the N-A2.2 variants with NIR which were tested once. “>>” means higher than 1000-fold. **D**) Crystallographic structure of the 6-helix bundle of the SARS-CoV-2 spike glycoprotein (PDB: 7tik [44]). HR1 is colored in grey and HR2 is colored in firebrick red. HR1 AA substitutions are annotated on the respective position in the 3D crystal structure and colored in grey. HR2 AA substitutions are annotated on the respective 3D crystal structure and colored in red. A black line spanning from residue D1168 to the utmost bottom of the 3D structure indicates the binding region of RQ-01. RQ-01 reaches residue K1205 of the spike, but it is not resolved in this crystal. **E**) Representative mWasabi kinetic curves of VSV-S-M^pro^ and RQ-A10.1, RQ-D4.2, RQ-D6.2 and RQ-D6.4 variants. Data is presented as mean ± SEM of 4 technical replicates. **F**) Bar plot of the TM_50_ values obtained from mWasabi kinetic experiments. Data is presented as mean ± SD of 2 independent experiments.

### VSV-S-M^pro^ RQ-A10.1 and RQ-D6.4 variants display faster mWasabi signal growth compared to the originating virus and other variants

Contrary to our expectations, two mutants generated by passage with RQ-01 also displayed a decreased susceptibility towards NIR (**Figure 3B**). As mutations can have an impact on virus kinetics, we measured routinely the untreated wells of every dose-response experiment at 12-hour intervals up to 96 hours after infection. We found that the apparent cross-resistant viruses produced more GFP earlier, indicating faster replication (**Figure 3E**). Therefore, the time to reach half of the maximal signal (TM_50_) of these two viruses was lower compared to another mutant and the parental virus (**Figure 3F**). In a standard multi-step replication kinetic, we confirmed that the same two mutants replicated faster and to higher titers than other mutants and the parental virus (**Supplemental Figure 2**).

### Passaged viruses adapt to the cell culture

We observed that after passaging against a selective pressure, some mutants appeared to replicate faster than others, thus displaying a mild ‘pseudo-resistance’. To assess whether such faster replicating viruses are adapting to cell culture, we passaged wild-type VSV-S-M^pro^ in the absence of any inhibitor. Indeed, some viruses arising from that passaging also displayed faster kinetics (**Supplemental Figure 3A,B**) and decreased susceptibility to the inhibitor NIR (**Supplemental Figure 3C,D**). However, compared to hallmark E166 NIR-resistant substitutions with an IC_50_ increase of >1000 fold, these cell culture-adapted viruses displayed milder phenotypes against NIR, the highest two being 37-fold (RQ-A10.1, **Figure 3C**) in a RQ-01 selected mutant and 19.3-fold in a virus passaged without inhibitor selection pressure (PWOI-A3.1, **Supplemental Figure 3E**).

### An orthogonal VSV pseudotype assay confirms spike HR1/HR2 mutants’ phenotypes

To confirm our results in a gold-standard pseudotype assay, we introduced different mutations into the HR1 and/or HR2 domains by point-mutating spike expression plasmids via PCR (**Supplemental Table 1**). Spike protein variants were then expressed by transfecting 293T cells with these expression plasmids. Upon infection of transiently transfected 293T cells with a VSV variant lacking the glycoprotein gene (VSV-ΔG), we produced VSV pseudotyped particles that displayed the spike protein on the surface instead of the native VSV-G. This allowed us to look at the unique contribution of HR1 and/or HR2 substitutions to the decreased susceptibility to RQ-01. With this approach, we could also test the selected mutations in the context of the natural XBB.1.5 furin cleavage site, which was mutated in our dual-chimeric VSV-S-M^pro^. We thus produced VSVs with spike HR1/HR2 variants that had shown reduced susceptibility to the fusion inhibitor in dose-response experiments with live viruses. We also included some variants of which we did not or could not purify the replicating VSV-S-M^pro^ version (e.g. HR2 single mutants R1185C and E1188D). We tested the first set of HR1/HR2 variants (**Figure 4A**), which included A944T, A944V, N953K, R1185C and N953K/R1185C. These variants arose from selection experiments with the parental VSV-S-M^pro^. We assessed a second set of variants (**Figure 4B**) that emerged from selection experiments with the N-A2.2 variant (harbouring the substitution E166V in M^pro^): T941K, Q949K, N953K, N953T, N955H, E1188D and N953H/E1188D. All these VSV-S-M^pro^_E166V variants included the parental spike harbouring an additional substitution – K1157N – as this variant appeared during passaging with the protease inhibitor NIR, while no selective pressure was being exerted on the spike. In **Figure 4C,D** we show the IC_50_ fold-changes of each spike variant compared to the unmutated parental spike to visualize the decrease in susceptibility to RQ-01.

**Figure 4.**
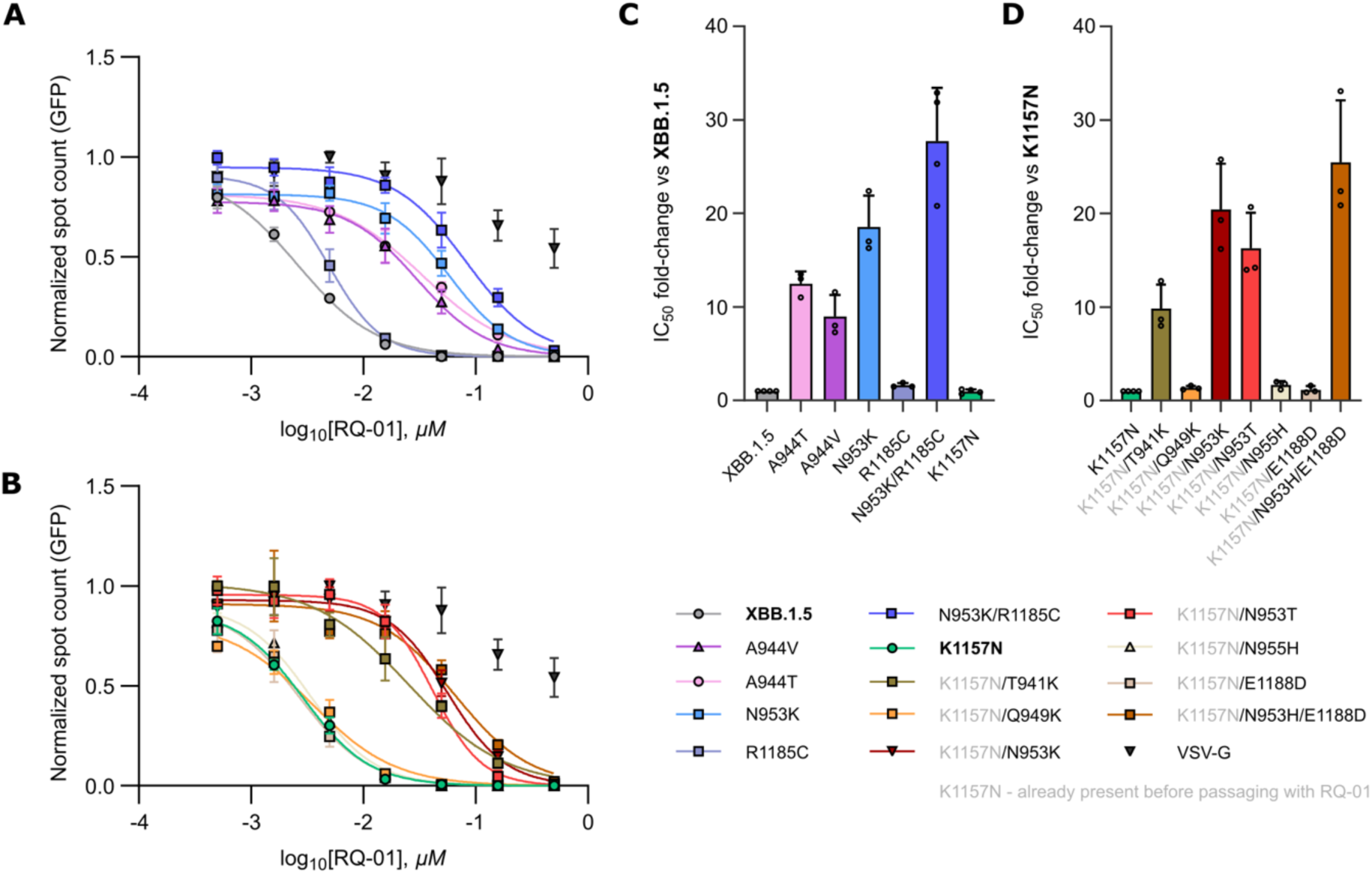
VSV pseudotype assay confirms spike HR1/HR2 mutant phenotypes. **A**) Representative RQ-01 dose-response assays of VSV spike pseudotypes harboring A944V, A944T, N953K, N953K/R1185C, R1185C, selected with VSV-S-M^pro^ vs unmutated spike (XBB.1.5 strain). Data is presented as mean ± SD of 3 technical replicates per condition. **B**) Representative RQ-01 dose-response assays of VSV spike pseudotypes harboring T941K, N953T, Q949K, N953K, N953H/E1188D, N955H vs spike-K1157N (original VSV-S-M^pro^_E166V spike variant where these substitutions arose from). Data is presented as mean ± SD of 3 technical replicates per condition. **C**) Bar plot of RQ-01 IC_50_ fold-change values for A944V, A944T, N953K, N953K/R1185C, R1185C vs unmutated spike (XBB.1.5 strain). Data is presented as mean ± SD of 3 or 4 independent experiments. **D**) Bar plot of RQ-01 IC_50_ fold-change values for T941K, N953T, Q949K, N953K, N953H/E1188D, N955H vs spike-K1157N (original spike variant where these substitutions arose from). Data is presented as mean ± SD of 3 or 4 independent experiments. In all the experiments, we included negative inhibition control by using VSV pseudotyped with its own glycoprotein G.

We observed that these results aligned well with our previous experiments and gave additional insights on the contribution of each variant. For example, the single variant N953K displays an IC_50_ fold-change of nearly 20, while R1185C did not affect the susceptibility of spike to RQ-01. The double variant N953K/R1185C was less susceptible to the inhibitor compared to N953K alone.

### Amino acid substitution selected in VSV-S-M^pro^ are present in GISAID deposited isolates

To investigate whether the amino acid substitutions selected in VSV-S-M^pro^ are compatible with SARS-CoV-2 replication, we used GISAID’s EpiCoV server to assess the number of entries for each substitution in actual virus isolates [45–47]. We found that some substitutions are quite frequent, such as L176F (29942 entries) and F186L (36302 entries), over all SARS-CoV-2 variants. The most frequent substitution in XBB.1.5 SARS-CoV-2 isolates was A348T (1149 entries). Substitutions in HR1 and HR2 domains were rare (**Supplemental Table 2**).

### ThermoMPNN analysis indicates that spike HR1 and HR2 variants are tolerated and do not strongly affect post-fusion stability

To account for the potential role of the experimentally tested spike variants to influence the stability of the post-fusion conformation, we carried out a two-state ThermoMPNN analysis, predicting the impact of the variants on the overall stability of the protein (**Figure 5A,B**). Several before experimentally tested mutations displayed negative scores, such as N955H, K968E in HR1 and R1185C in HR2 shown in the bar plots in **Figure 5C** and **5D**, indicating a preference to the post-fusion conformation. Other experimentally tested mutations were also predicted to slightly stabilize this conformation, which may simply reflect a tolerance to the post-fusion conformation. Three mutations (A944V, E988D for HR1 and P1162Q for HR2) were predicted to have a post-fusion destabilizing effect. As the ThermoMPNN predictions are sensitive to the choice of input backbone, the scores, particularly for residues proximal to the terminal regions of the protein, should be interpreted with caution. This applies to the E988D and P1162Q mutation, as they are the C-terminal residue of HR1 and the fourth residue from the N-terminal region in HR2 of the experimentally determined structure, respectively (PDB accession: 2FXP [48]).

**Figure 5.**
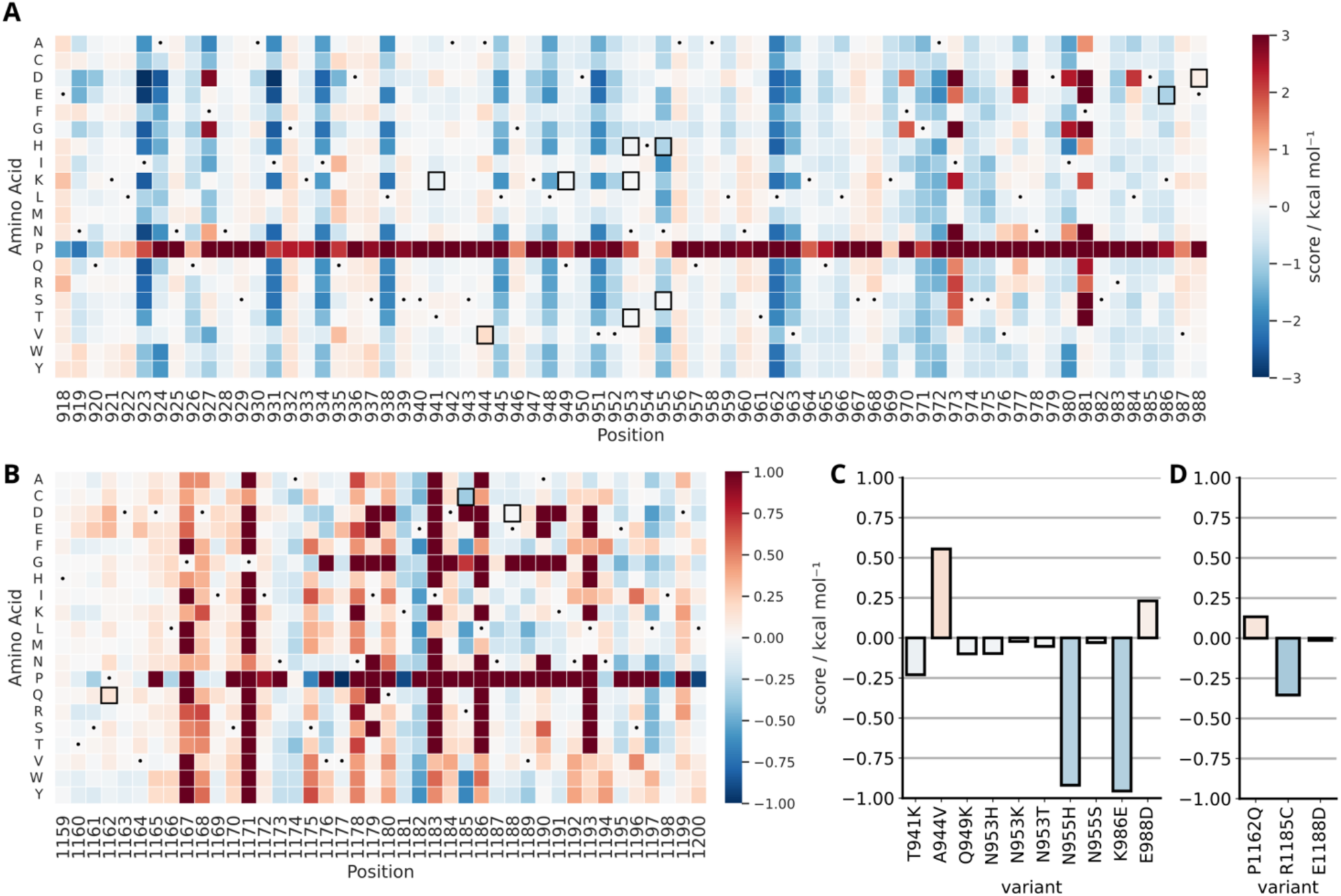
ThermoMPNN analysis indicates that spike HR1 and HR2 variants are tolerated and do not strongly affect post-fusion stability. The color scale represents the predicted change in stability of the post-fusion conformation relative to the pre-fusion conformation, ranging from post-fusion stabilizing (blue) to post-fusion destabilizing (red), calculated in kcal·mol⁻¹. **A**) Heat map of the HR1 region showing per-residue, per-substitution stability scores for all possible amino acid substitutions. **B**) Heat map of the HR2 region using the same scale. Black dots indicate the wild-type amino acid at each position. Empty black squares indicate experimentally investigated variants (HR1 and HR2). Bar plots of the stability scores for experimentally investigated variants in HR1 (**C**) and HR2 (**D**).

### Classical molecular dynamics simulations provide detailed information on native HR1-HR2 and HR1-RQ-01 interaction pattern

The SARS-CoV-2 six-helix bundle (6-HB) comprises six α-helices in a complex formed by the heptad repeat elements HR1 and HR2. In this structure, three central HR1 helices form a trimeric coiled coil, while three HR2 helices pack in an antiparallel orientation into the hydrophobic grooves of HR1 [49,50]. Overall, the 6-HB structure is characterized by a complex network of hydrophobic, hydrogen-bonding, and ionic interactions [51]. Hydrophobic interactions are mainly present in the helical fusion core region and are the primary driver of the association of these domains, involving residues between V1164 and I1198 in the HR2 domain [50]. In addition to hydrophobic packing, electrostatic and polar interactions contribute to the structural stability [52]. Two conserved inter-helical salt bridges, D936-R1185 and K947-E1182, are essential for stabilizing HR1-HR2 packing, thus enabling membrane fusion.

Notably, the mutation-induced disruption of the D936-R1185 interaction can destabilize the complex, resulting in a loss of cell-cell fusion activity [51]. Hydrogen bonds are mediated by HR1 sidechains interacting with both backbone and sidechains of HR2. In the reference structure (PDB ID: 7TIK [44]), multiple hydrogen bond interactions are observed (**Figure 6A**). Among these, Q949-I1179, S943-E1182, and N960-A1174 hydrogen-bond interactions are widely discussed in the literature [49,53,54]. In contrast, other interactions such as N928-I1198 [55], S943-E1182 [52] are less frequently reported, while additional interactions, including N953-V1177, S940-E1182, Q935-N1194/A1190, and Q949-Q1180 [54] are not present in this structure but appear in other deposited structures.

**Figure 6.**
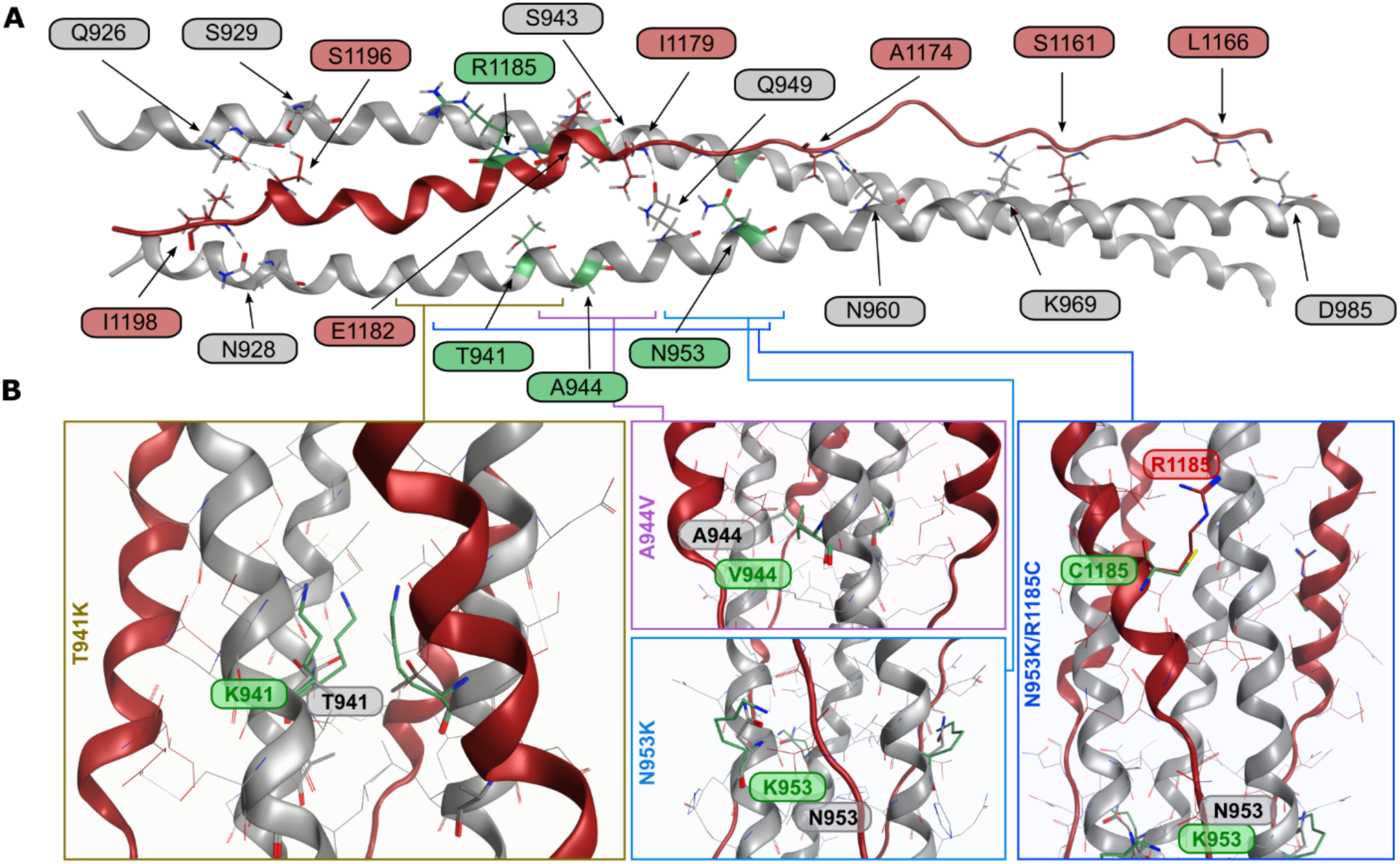
Structure of the HR1/HR2 bundle: interaction, important residues and close-up of selected variants. **A**) Hydrogen bonds between HR1 monomers (grey) and HR2 monomer (fire brick) in the 7TIK [44] structure (6-HB). AA engaging in hydrogen bonds between HR1 and HR2 are labeled in grey and fire brick, respectively. **B**) Superimpositions of the spike structures with the substitutions T941K, A944V, N953K and N953K/R1185C (in place) and the structure of the parental spike. The original residues are labeled in grey and substituted in green. Their positioning in the HR1/HR2 coiled structure is indicated by highlighting them on the structure shown in panel **A**.

To investigate the dynamic behavior of the natural or ‘non-stapled’ 6-helix complex, we performed classical molecular dynamics (MD) simulations of 100 ns on the non-stapled HR1/HR2 (nstap-HR1/HR2; 6-HB) and the stapled HR1/HR2 (stap-HR1/HR2; (RQ-01)-(5-HB)) complexes. The stapled 5-helix bundle (5-HB) complex was set up to leave one binding site empty for RQ-01 binding. Both complexes remained sufficiently stable throughout the simulations, with average RMSD values – calculated across three replicas – of 1.78 Å for the nstap-HR1/HR2 and 2.50 Å for the stap-HR1/HR2. Notably, the alpha-helix region of RQ-01 remained particularly stable, with average RMSD values of 1,23 Å for the nstap-HR1/HR2 and 1,12 Å for the stap-HR1/HR2. As illustrated in the per-residue interaction energy plots (**Supplemental Figures 4,5**), the overall interaction pattern is comparable between the two complexes. Key hydrogen-bond interactions including Q949-I1179, N960-A1174, and N928-I1198 were retained throughout the simulations, while others such as N953–V1177, Q935-N1194/A1190, and Q949-Q1180, were formed during the simulation and remained stable. Several residues exhibited a more dynamic behavior, interacting with multiple amino acids. For example, E1182 primarily forms a salt bridge with K947, but can also interact transiently with both S943 and S940. Similarly, R1185 is involved in a salt bridge with D936, and occasionally interacts with E1182, disrupting the E1182-K947 salt bridge. In **Figure 6B**, we provided a structural representation of the spike variants investigated through Thermal Titration Molecular Dynamics (TTMD) simulations, outlined in the following section.

### Thermal Titration Molecular Dynamics (TTMD) simulations reveal the impact of selected spike HR1/HR2 variants on the stability and protein-peptide interaction with RQ-01

To examine the stability of the (RQ-01)-(5-HB) complex while accounting for system flexibility, we carried out TTMD simulations. This technique ultimately returns a MS coefficient, which describes the stability of the complex between RQ-01 and the 5-HB under a stepwise increase of thermal stress. A lower MS coefficient means a more stable complex, whereas a higher MS means a less stable complex. This approach was previously applied to study different variants of the SARS-CoV-2 M^pro^ compared to the wild type (Wuhan-1 strain) [20]. In the current work, the unmutated (RQ-01)-(5-HB) complex stap-HR1/HR2 was compared to different complexes involving single and double mutations of the (RQ-01)-(5-HB / A944V, T941K, N953K, N953K/R1185C) complex. TTMD analyses were carried out, and the resulting MS coefficients were used to rank the variants from the most to the least stable: N953K > N953K/R1185C > A944V > T941K (**Supplemental Table 3**). We observed that only A944V and T941K were less stable than the unmutated complex.

To therefore refine our approach, we reanalyzed the molecular dynamics simulations by “splitting” RQ-01/HR1 complex in two portions and rescored the simulation according to where the mutations occurred. First, the alpha-helical RQ-01 part for T941K and A944V (rescoring alpha-helix **Supplemental Figure 6A-C**) and second, the C-terminal RQ-01 part for N953K and N953K/R1185C (rescoring C-terminal portion, **Supplemental Figure 6D-E**).

We thus recalculated the MS score for T941K and A944V considering only the alpha helical portion of HR2 (I1179-S1196), since it is directly in contact with these residues. This rescoring improved the MS score difference between the T941K and A944V stap-HR1/HR2 mutants and the unmutated stap-HR1/HR2, emphasizing their higher predicted instability (**Supplemental Table 4**). After rescoring, T941K had a higher mean MS value (0.00358) than the unmutated stap-HR1/HR2 complex (0.0029). We observed a temperature-dependent loss of native interactions in the titration timeline plot for the T941K variant (**Supplemental Figure 6B**) and high instability (**Supplemental Video 1**). In addition, a root mean square fluctuation (RMSF) plot showed the increased flexibility of K941 (**Supplemental Figure 7**). The rescoring of A944V led to a mean MS score of 0.00311 (**Supplemental Figure 6C**) (vs 0.0029 of the unmutated stap-HR1/HR2), indicating increased predicted instability (**Supplemental Video 2**).

N953K alone had a lower score than the full stapled complex (stap-HR1/HR2) (**Supplemental Table 3**). As this substitution occurred together with R1185C, a residue which interacts with the C-terminal part of RQ-01, we rescored N953K alone and N953K/R1185C together against the C-terminal portion of RQ-01 (D1168-N1178). The MS scores of both variants indicate a slightly higher predicted complex instability for the mutants compared to the wild type, with mean values of 0.00550 for N953K and 0.00615 for N953K/R1185C, versus 0.0053 for the unmutated stap-HR2 (**Supplemental Table 5**). We observed in the simulation that N953K disrupts a hydrogen bond formed by N953, whereas the contribution of R1185C to the complex instability is less obvious and probably mostly relevant in concert with N953K (**Supplemental Videos 3,4**).

## Discussion

Antivirals are life-saving medicines. However, viruses can evolve quickly, and antivirals are under constant threat to become ineffective. When resistance occurs in the clinical setting, ideally the mechanism and resistance spectrum of the virus and drug(s) are already known, so health care professionals can react quickly. In such a scenario, the drug regimen can be switched to successfully treating the resistant virus. Furthermore, some chronic virus infections (i.e. HIV-1) require a priori treatment with several drugs to prevent resistance. Therefore, viral resistance and cross-resistance studies before the licensing of a drug or in parallel to its real-life use are essential, but also controversial due to biosafety concerns of GOF research. To address this conflict, we developed a safe alternative in form of a dual-chimeric vesicular stomatitis virus, VSV-S-M^pro^. With this virus we were able to study the development of resistance against two different classes of coronavirus inhibitors, namely main protease (M^pro^) and spike fusion inhibitors. Thus, we could test remaining susceptibility and cross-resistance of mutants against these two inhibitor classes.

In this study we first selected mutants against NIR with VSV-S-M^pro^. We then tested the susceptibility of hallmark M^pro^ E166 mutants [39–41] against NIR in comparison to an anti-XBB.1.5 spike serum and the fusion inhibitor RQ-01 [37]. RQ-01 is a peptide fusion inhibitor that mimics the HR2 domain of spike and the peptide is linked to a membrane anchor. Unsurprisingly, we observed that M^pro^ mutants were resistant to NIR, but not the XBB.1.5 spike antiserum and RQ-01. We then selected RQ-01 mutants with parental VSV-S-M^pro^ and VSV-S-M^pro^-E166V. As expected, RQ-01-selected VSV-S-M^pro^ mutants had reduced susceptibility to the fusion inhibitor, but in general not NIR. Curiously, there were some few RQ-01-selected mutants that also showed a mild resistance phenotype against NIR. In kinetic experiments, those mutants replicated faster than others and their parental virus. To test whether cell culture adaptations could happen in VSV-S-M^pro^ (observed in SARS-CoV-2 [56]), we passaged virus without any inhibitor. Indeed, some such passaged viruses displayed mild resistance phenotypes against NIR, albeit by far lower IC_50_ fold changes than hallmark mutants such as E166K/V. Lastly, RQ-01-selected mutants of VSV-S-M^pro^-E166V had reduced susceptibility to the fusion inhibitor and pronounced resistance to NIR at the same time.

During cloning and rescue of the dual-chimeric VSV-S-M^pro^ and subsequently passaging with the M^pro^ inhibitor NIR, we detected the amino acid substitutions R682W and K1157N, respectively. These mutations emerged independently from the selection pressure stimulus exerted by RQ-01. R682W most likely appeared due to an adaptation to the Vero ↑ACE2 ↑TMPRSS2 culture [38]. K1157N, a residue in the proximity of the HR2 domain, emerged in sample N-A2.2 during selection pressure experiments with the protease inhibitor NIR and therefore was ubiquitously present in all the viral progeny of N-A2.2 that was selected with RQ-01. Moreover, most of the tested RQ-01-selected viruses had multiple amino acid substitutions along the whole spike sequence and not only in HR1 or HR2 domains. Since we could not test the contribution of all mutations, we focused on a few HR1 and/or HR2 variants by using pseudotyped VSV-Spike HR1/HR2 variants. Most HR1 variants exhibited decreased susceptibility to RQ-01, while HR2 variants did not. This result could be explained by the fact that some mutations can happen independently from the selective pressure put by the HR2-decoy RQ-01. One example is E1188D (HR2 domain), which is a substitution described by another group looking at cell culture adaptations of SARS-CoV-2 after passaging several times on Vero E6 cells [38]. Nevertheless, the combination of a HR1 and a HR2 – such as in N953K/R1185C – substitution caused a decrease of RQ-01 activity. We could also confirm that the spike variant K1157N was as susceptible as the parental spike (XBB.1.5) to RQ-01, whereas VSV-G was not susceptible, as expected. We successfully confirmed our live-virus results by employing an orthogonal, gold-standard method, a VSV-pseudotype assay. This type of assay makes the majority of the viral particle uniform between variants, except the spike. Therefore, different virus behavior can be attributed specifically to changes in spike.

In a protein stability analysis using ThermoMPNN, we observed that different substitutions affect the post-fusion conformation stability either positively or negatively. The energetic contribution was mostly negligible, suggesting that these mutations were well tolerated but may still partially influence the final conformation.

In molecular dynamics simulations we could investigate molecular interactions in more detail through TTMD analyses. Collectively, we observed a variety of effects, based on the position and type of amino acid substitution. For example, our simulations suggest that T941K substantially altered the wild-type binding mode, providing a mechanistic explanation for its impact on the decreased efficacy of RQ-01. The substitution A944V exerts its effect by a steric clash in the inner part of the 3-helix bundle formed by the three HR1 monomers. On the other hand, the substitution N953K results in enhanced flexibility of the local environment and of the close residues. However, these results may be influenced by the intrinsic flexibility of the tail portion, further amplified by high temperatures. Finally, we speculate that the simultaneous presence of the R1185C further destabilizes the (RQ-01)-(5HB) complex by weakening local cohesion between HR1 and HR2 due to the loss of the salt bridge between R1185 and D936, as already reported for the D936Y mutation [51].

Although the extent of perturbations caused by each substitution on the RQ-01/HR1 interface remains uncertain, we hypothesize that the destabilization of the HR1-HR2 interaction could be tolerated by the membrane-associated HR1 and HR2 monomers, whereas RQ-01 may be more vulnerable given its reliance on a lipid anchor. Alternatively, allosteric effects, not observed in these simulations, or altered monomer association may be involved, requiring further investigation to fully elucidate the role of these mutations.

In conclusion, we demonstrated that dual-chimeric VSV-S-M^pro^ is a useful surrogate tool allowing safer GOF studies that with more dangerous viruses would not be possible, undergo scrutiny or require high-containment infrastructure (BSL-3 or -4 labs). With this study, we aim to highlight the importance of this essential branch of virology, but at the same time propose the development of safer options to reduce drastically the risks associated with performing GOF experiments.

## Methods

### Cell lines

293T (ATCC), 293T A4 (expressing ACE2) and Vero cells (expressing ACE2 and TMPRSS2) cells were cultured in Dulbecco’s Modified Eagle Medium (DMEM) supplemented with 10% FCS, 2% glutamine, 1% sodium pyruvate, 1% non-essential amino acids (Gibco) and 1% Penicillin/Streptomicin.

### Cloning strategies

For the plasmid encoding the genome of VSV-S-M^pro^ (VSV-PmWasabi-Spike-M^pro^), the cloning strategy was carried out in three steps.

*First*, we performed restriction digestion of VSV-M^pro^ plasmid [17] with MluI and BbvCI to remove the native glycoprotein of VSV (VSV-G). We ran a PCR on cDNA obtained from several swab-derived SARS-CoV-2 XBB.1.5 isolates, amplifying XBB.1.5 spike sequence with a short 18 amino acid truncation at the C-terminus with primers “VSV-Spike_XBB.1.5 for” and “VSV-Spike_truncated_XBB.1.5 rev” (**Table S1**), as it improves incorporation of the spike into the virion [57]. We used the PCR with the highest yield and assembled it with the digested VSV-M^pro^ plasmid in a Gibson assembly reaction, generating a plasmid encoding the first version of the double chimeric VSV, lacking the reporter gene mWasabi (VSV-Spike_truncated_XBB.1.5-M^pro^). We then rescued a replicating VSV variant, plaque purified it in Vero ↑ACE2 ↑TMPRSS2, sequenced the spike protein and detected a mutation in the furin cleavage leading to the substitution R682W (^681^H**R**RARS^686^ to ^681^H**W**RARS^686^). We used this spike sequence for the final VSV-S-M^pro^ plasmid.

*Second*, we cloned the marker gene mWasabi in VSV-P to seamlessly visualize viral replication as previously shown [58] with primers “N-for” and “P-rev” (**Table S1**) from the sequence of “VSV-PmWasabi-L-mCherry”. To that end, we carried out another restriction digestion of the VSV-M^pro^ plasmid [17] with BstZ17l and XbaI to remove a fragment stretching from N to P. We then replaced the missing fragment with the PCR product N-PmWasabi described above via Gibson assembly, generating VSV-PmWasabi-M^pro^.

*Third*, we digested the plasmid encoding VSV-PmWasabi-M^pro^ with MluI and BbvCI to remove the native glycoprotein of VSV (VSV-G). As above, we replaced G with the spike glycoprotein using Gibson assembly. The spike sequence was derived from VSV-Spike_truncated_XBB.1.5-M^pro^ with the furin cleavage site mutation via a PCR with primers “VSV-Spike_XBB.1.5 for” and “VSV-Spike_truncated_XBB.1.5 rev”.

### Gibson assembly

We carried out Gibson assembly cloning [59] following manufacturers’ instructions (NEBuilder HiFi DNA Assembly Master Mix, NEB) to obtain circularized plasmids from a vector plasmid and one or two inserts/fragments. We digested the starting vector beforehand with restriction endonucleases. We added inserts and vectors together in a molar ration of 1:3 in a final volume of 10 or 20 μL. As preparation for Gibson assembly, the inserts had to be PCR-amplified with primers containing overhangs (from 20 to 35 bp) complementary to the 5’ and 3’ ends of the digested vector. We incubated the reactions at 50°C for four hours in a thermocycler.

### Viral rescue

One day before rescue, we seeded 2-3×10^6^ 293T cells in 10-cm dishes in 8 mL of supplemented DMEM. The next day, the rescue was performed when cells were between 50 and 70% confluent. We prepared a master mix of the plasmid encoding the genome of VSV-S-M^pro^ with helper plasmids, as described previously [60], into a 1.5 mL microcentrifuge tube and filled up with sterile, nuclease-free water to 450 μL. Then, we added 50 μL of 2.5 M calcium chloride (CaCl_2_) and vortexed. We transferred the mix to a 50 mL centrifuge tube containing 500 μL of HEPES 2X buffer in a dropwise fashion under constant vortexing. We incubated the mix for 20 minutes at room temperature to allow the formation of calcium phosphate-DNA precipitates. In the meantime, we premixed 2 mL of supplemented DMEM with 10 μL of 25 mM chloroquine and carefully added the solution to the seeded 293T cells to reach a final concentration of 25 μM. After the 20 minutes incubation time we transferred the mix to the 293T cells, which we incubated at 37°C and 5% CO_2_ in a humidified incubator. Chloroquine is cytotoxic, therefore in a timeframe between 6 to 16 hours post transfection we replaced the chloroquine-containing medium with fresh cell culture medium. Once cytopathic effect was visible, we collected the supernatant, filtered it through 45 μM filters, aliquoted it and stored at -80°C for subsequent plaque purification.

During the rescue, we observed the VSV-S-M^pro^ was replicating poorly and maintaining virus over time was difficult. We hypothesized that its highly altered genome attenuated viral replication. Indeed, it had been described that genome alterations in VSV can lead to temperature sensitivity [58,61,62]. Therefore, we lowered the temperature for culture from 37° to 34°C after the initial rescue at 37°C. As this drastically improved replication, we performed all following experiments in a humidified incubator at 34°C and 5% CO_2_.

### Plaque purification of VSV-S-M^pro^ after rescue

We seeded 2-3×10^5^ Vero↑ACE2 ↑TMPRSS2 cells/well in 6-well plates containing 2 mL of cell culture medium each well one day prior infection. Before removing the 2 mL of medium, we prepared serial dilutions of the virus to be plaque-purified (1:10 dilution steps). We removed medium from the cells using a vacuum pump, added one virus dilution for each well and incubated at 34°C and 5% CO_2_ in a humidified incubator. As the incubation step takes 1.5 to 2 hours, we prepared a 1:1 mixture of low melting plaque agarose with culture medium and kept it pre-warmed at 37°C in a water bath. After 1.5-2 hours of incubation, we aspirated the virus-containing supernatant from each well with a vacuum pump. Immediately after that, we added 2 mL of 1:1 mixture of pre-warmed plaque agarose and culture medium on the wells and left the plates outside the incubator for 5 to 10 minutes to allow the plaque agarose to solidify. Then, we placed the plates at 34°C and 5% CO_2_ in a humidified incubator. We checked the formation of green plaques (mWasabi+) daily until they became large yet distant enough to be picked singularly. Finally, we infected 293T A4 (↑ACE2) cells seeded in a 6-well plate one day prior with the picked viral plaques, each one in a separate well of the 6-well plate. We let the virus replicate for three to four days, collected the supernatant, prepared aliquots and stored them at -80°C until further use.

### Determination of infectious virus titer (TCID_50_/mL)

To determine the viral titer of our stocks, we carried out tissue culture infective dose fifty (TCID_50_/mL) assays, which quantify the amount of infectious virus in a viral sample. We seeded 1-2×10^3^ cells/well in 100 μL in 96-well plates one day prior to infection. On the day of infection, we serially diluted the virus in a logarithmic manner (base 10) down to 10^-9^ and then added 100 μL of each virus dilution to 100 μL of the cells seeded one day before. We incubated the plates at 34°C and 5% CO_2_ in a humidified incubator for 5-6 days, after which we counted the number of mWasabi+ wells, which indicate a successful infection. We then calculated the virus titer(s) using the Spearman and Kaerber formula [63].

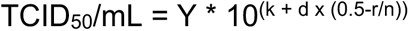

Y: Extrapolating factor to get TCID_50_/mL, Y=10 when 100 μL dilution/well
k: positive exponent of the highest dilution tested
d: spacing between dilutions, d=1 for log and d=0.5 for half log
r: total number of wells that were infected but CPE/mWasabi negative
n: number of wells per dilution

### VSV-S-M^pro^ production for experimental use

We produced stocks of the original VSV-S-M^pro^ using 293T A4 (↑ACE2) cells. We seeded 293T A4 cells in 15 cm plates at a density of 11×10^6^ cells in 20 mL of medium (fully supplemented DMEM). One day after seeding, we infected cells with VSV-S-M^pro^ at 0.01 or 0.02 MOI and incubated at 34°C and 5% CO_2_ in a humidified incubator until mWasabi signal and clear cytopathic effect was visible. After 48-60 hours post infection, we collected the supernatant and filtered it with 45 μm filters in a 50 mL centrifuge tube. Afterwards, we aliquoted the filtered supernatant in 1.5 ml screw-cap tubes and placed them at -80°C for further titration and use.

### Dose-response assays with live VSV-S-M^pro^

For dose-response experiments, we seeded 1×10^4^ 293T A4 (↑ACE2) cells/well in 50 μL of medium in 96-well plates. On the same day, we prepared viral dilutions in cell culture medium to a MOI of 0.01 in each well of the 96-well assay plates (with the only exception of dose-responses in **Figure 2C-D**, where the MOI was 0.1). We then distributed the virus plus medium into one row per virus variant of a 96 deep-well plate. After adding the appropriate volume of stock solution of inhibitor to the leftmost wells (first column of the plate), we started a three-fold serial dilution transferring the inhibitor dilution from one well to the next one from left to right until the 11th column of the deep-well plate. The 12th column was left without inhibitor and served as untreated control. For each virus variant, we calculated the required volume of virus medium and inhibitor needed for two to four technical replicates. We further added extra volume to account for medium sticking to well plate surfaces. After that, we transferred 50 μL of the prepared serial dilutions to the seeded cells. We gently hit the plates four to five times on each of the four sides and checked whether cells were homogeneously distributed. We then placed the plates in a wet chamber and put them in a humidified incubator at 34°C and 5% CO_2_.

For the dose-responses with anti-XBB.1.5 serum, we started the three-fold serial dilution with prediluted serum (1:20) and proceeded as described above.

Starting from 12 hours post infection, we read out the mWasabi signal at approximately 12-hour intervals with an ELISpot analyser (Series 6 ULTRA-V) using the FluoroSpot X Suite provided in the proprietary software package from C.T.L. (Cellular Technology Limited). The sampling every 12 hours is described in more detail in the paragraph “mWasabi signal growth kinetics”. Regarding the dose-response read out, we generally captured the mWasabi signal at 48-60 hours post infection, but in some cases, we read out also at later timepoints. After 48-60 hours post infection, we loaded the plates (without removing the medium) in the analyser to scan each well of the 96-well plates and sample the mWasabi spots within the wells. At the counting step, we set the counting area to 70% to exclude the optically distorted outer area of the wells due to the medium as well as reflection on the walls of the wells. The Basic Count program automatically generates an excel spreadsheet containing the raw data. The raw data analysis is described later in this section.

### Serial passage of VSV-S-M^pro^ with NIR and ENS to select M^pro^ mutants

For the first passage of a series of ten passages with the M^pro^ inhibitors NIR and ENS, we seeded 293T A4 (↑ACE2) at a density of 5×10^3^ cells/well in 100 μL of medium in a 96-well plate. We divided the 96-well plate in half (on the long side), and assigned one half to NIR and one to ENS. At the same day of seeding, we premixed VSV-S-M^pro^ at an MOI of 0.01 and NIR or ENS to a final concentration of 0.5 μM after the addition of the virus and compound mix (100 μL) to the cells, reaching 200 μL final volume in each well. We placed the plate at 34°C and 5% CO_2_ in a humidified incubator and incubated for 3-4 days. For the subsequent passages, we seeded 5×10^3^ 293T A4 cells/well in 80 μL of medium and then transferred 20 μL of supernatant from the previous passage to the freshly seeded cells. Subsequently, we added 100 μL of inhibitor, thoroughly pipetting up and down to ensure proper mixing of the compounds and homogeneous distribution of the cells. We incubated the plates 3-4 days for each passage at 34°C and 5% CO_2_ in a humidified incubator. Up to passage 7, we doubled the concentration of inhibitor each time, reaching up to 32 μM of either NIR or ENS. For passages 7, 8 and 9 we kept a final concentration of 32 μM to avoid losing too many positive wells. For passage 10 we increased the concentration up to 64 μM. The sequence of inhibitor concentrations from passage 1 to passage 10 was 0.5, 1, 2, 4, 8, 16, 32, 32, 32, 64 μM. The supernatants of mWasabi+ wells were used for sequencing experiments (100 μL), for plaque purification (50 μL) and production of selected variants.

### Serial passage of VSV-S-M^pro^ with RQ-01 to select spike mutants

For the serial passage with the fusion inhibitor RQ-01, we proceeded as described above. We infected one full 96-well plate for each passaged virus (parental VSV-S-M^pro^ and the VSV-S-M^pro^-E166V/N-A2.2 obtained after passaging with NIR). We started with a concentration of 1 nM of RQ-01 and doubled it up until passage 8, reaching a concentration of 128 nM. For passages 8, 9 and 10 we kept a final concentration of 128 nM to avoid losing too many infected wells. The sequence of inhibitor concentrations from passage 1 to passage 10 was 1, 2, 4, 8, 16, 32, 64, 128, 128, 128 nM. The supernatants of mWasabi+ wells were used for sequencing experiments (100 μL), plaque purification (50 μL) and production of selected variants.

### Plaque purification and production of VSV-S-M^pro^ variants after serial passaging/selection experiments

To plaque purify the selected virus variants, we seeded 2.5-5×10^4^ 293T A4 (↑ACE2) cells/well in 24-well plates containing 500 μL of cell culture medium each well one day prior infection. Before removing the supernatant, we prepared serial 1:10 dilutions of the virus to be purified. We then proceeded as described above and added 250 μL of a 1:1 mixture of pre-warmed plaque agarose and culture medium to the wells and left the plates outside the incubator for 5 to 10 minutes to allow the plaque agarose to solidify. After 3-4 days, we marked the plaque locations with a marker on the outer bottom of the plate. In the plaque picking step, we pierced through the plaque agarose layer at the marked location with a pipet tip to collect the virus and pipetted the retrieved material up and down in a 1.5 microcentrifuge tube containing 1 mL of medium. Subsequently, we infected 293T A4 cells seeded at a density of 1.5-2×10^5^ cells/well in a 6-well plate one day prior infection with the picked viral plaques. We let the virus replicate for 3-4 days, collected the supernatants, prepared aliquots and stored them at -80°C until further use. Finally, we infected 293T A4 (↑ACE2) cells seeded at a density of 1.5-2×10^5^ cells/well in a 6-well plate one day before infection with the viral stocks derived from the picked plaques. We incubated cells and viruses for 3-4 days until extensive cytopathic effect (CPE) and mWasabi signal were visible, collected the supernatants, filtered through a 45 μM filter, prepared aliquots and stored them at -80°C for experiments.

### mWasabi signal growth kinetics

When performing dose-response experiments, we left the last column of the 96 well/plate untreated. The virus (VSV-S-M^pro^ and related variants) can then grow without being affected by the inhibitor(s). In this setting, we can use the rate at which the mWasabi signal grows over time to monitor viral replication speed. We started reading out the mWasabi signal spot count at 12 hours post infection and then imaged every 12 hours. We sampled the mWasabi spot count up to 96 hours post infection, but in some cases, we stopped at either 84 or 72 hours as the mWasabi spot count number started to decrease due to virus-induced cell death. We scanned and counted the mWasabi spots with the same settings and parameters used for the read out of dose-response experiments with VSV-S-M^pro^.

### Standard multi-step growth curve of VSV-S-M^pro^ variants with TCID_50_/mL readout

We seeded 293T A4 (↑ACE2) cells at a density of 1×10^5^ cells/well in 24-well plates one day before infection. At the day of infection, we prepared viral dilutions at 0.01 MOI, infected the cells in duplicates for each condition and then incubated the plates in a humidified incubator at 34°C and 5% CO_2_. We collected supernatants in 1.5 mL screw-cap microcentrifuge tubes at the desired timepoints (0, 8, 25, 48 and 60 hours post infection) and stored them at -80°C until further use. We performed titration experiments for each condition, two duplicates for each timepoint, with TCID_50_/mL as read out.

### Viral RNA isolation, cDNA synthesis and PCRs for ONT sequencing

We isolated all viral RNA using the E.Z.N.A. Viral RNA Kit from Omega Bio-Tek. The maximum sample volume to be used in this kit is 150 μL. We used this amount, when possible, except when the available volume was insufficient, for example from a well of a 96-well plate, where we used only 50 or 100 μL instead. After the isolation of viral RNA, we proceeded with the synthesis of cDNA using the RevertAid RT Reverse Transcription Kit (ThermoFisher Scientific). Finally, we ran PCR reactions using the synthesized cDNA as a template to generate amplicons of the region of interest to be sequenced. Primer pairs (**Table S1)** were the following: “vsv-M_qPCR_for” plus “L N-terminal rev” or “VSV_Spike-seq_for” plus “VSV_Spike-seq_rev” for the spike protein and “cut1-for” plus “L N-terminal rev” or “VSV_1764bp_after-M_for” plus “L N-terminal rev”. We checked if–and how many–reactions worked after amplification of the target regions from viral cDNA by running 1-2 μL of PCR product on 1% agarose gels, stained with GelRed 3X for 30 minutes and visualized with a GelDoc-it system (UVP).

### ONT sequencing workflow

We prepared sequencing libraries using the Rapid Barcoding Kit 96 (SQK-RBK110.96) provided by ONT and followed the manufacturer’s protocol. Once a library was prepared, we stored it on ice until loading into the flow cell. Before loading, we primed a R.9.4.1 MinION flow cell following the manufacturer’s protocol. We then loaded the library into the flow cell through the Spot-On port and started sequencing through the proprietary software MinKNOW. Parameters of the run: kit selection (SQK-RBK110.96) > trim barcodes > Q score cutoff = 8 > raw data file format = .pod5.

After 24 to maximum 48 hours of sequencing, we manually stopped the run. We transformed the raw sequencing data from electrical signals (.pod5) to nucleotide bases (.fastq) using the basecaller Guppy (version 6.5.7) with the super-high accuracy model. Once we retrieved the nucleotide sequence of each read and for each sample/barcode, we merged the reads of each barcode (several .fastq files for each barcode) in one large file containing all the reads of that specific barcode. We then filtered raw reads to a PHRED quality score of ≥Q15 and length depending on the size of the PCR amplicon using SeqKit (version 2.4.0 [64], aligned to the reference sequence using minimap2 (version 2.22) [65], followed by sorting and indexing using SAMtools (version 1.13) [66]. SAMtools depth was used to check sufficient depth. To find SNVs, we used LoFreq (v2.1.3.1) for variant calling with—min-cov option set to depth = 300 (= 100 when a portion of the amplicon had less than 300 reads) [67]. This workflow generated several file formats from which we used the .vcf (variant calling fille) to visualize the mutations detected in the samples. ONT data analysis scripts are available in **supplemental File S1**.

### Post-processing of sequencing data after analysis

Once we obtained the .vcf files, we uploaded them into the software Geneious Prime and mapped them to a reference sequence. The software annotated/mapped the information in the .vcf file on the reference sequence and we were able to visualize the position of each nucleotide. From here, we exported the .csv file containing the information on barcode number, nucleotide change, position in sequence and frequency of this mutation and other parameters. After the export, we used an excel spreadsheet that automatically calculated the amino acid exchange and position in the protein sequence, returning the amino acid substitution format XNNNY (X = old AA; Ns = position number; Y = new AA). We checked these mutations manually for plausibility.

### Dose-response data analysis (IC_50_, IC_50_ fold-changes and TM_50_)

To obtain IC_50_ values from the dose-response data with live VSV-S-M^pro^, we used the excel spreadsheet generated after the Basic Count program. We normalized the spot count data of each dataset individually to the maximum average value and obtained a range of values between 0 and 1. We displayed the normalized data on the y-axis, against the inhibitor concentrations, plotted on the x-axis. After that, we ran a non-linear regression analysis using the equation “Sigmoidal, 4PL, X is concentration”, which is pre-set in the software GraphPad Prism. We added a constraint to the bottom value “BOTTOM=0” as we noticed that the curve fit returned unrealistic bottom value lower than 0. The analysis generates an additional sheet containing the inhibitory concentration fifty (IC_50_) value. We compared the IC_50_ values of the inhibitors used for dose responses assays (anti-XBB.1.5 serum, NIR, ENS, BOF or RQ-01) between VSV-S-M^pro^ variants and the parental/original virus to assess the relative change in efficacy of the inhibitors towards the mutant viruses.

For time-maximum fifty (TM_50_) calculations, we used the data sampled at each 12-hour interval for the mWasabi signal growth kinetics. After data collection, we normalized the spot count data as described above. We displayed the normalized data on the (y-axis) over time (x-axis). We performed non-linear regression analysis using the equation “Sigmoidal, 4PL, X is hours”, a renamed version of Sigmoidal, 4PL, X is concentration. We defined three constraints: HillSlope < 15; bottom value “BOTTOM=0”; top value “TOP=1”. Moreover, for fitting the curves, we excluded the data points after the highest value of each individual dataset to avoid the influence of the spot count decrease (cell death) at late timepoints to the TM_50_ value. From this analysis, we obtained the extrapolated values that we called time maximum fifty (TM_50_), the time needed to reach half of the maximum signal/the plateau.

### Generation of spike-pseudotyped VSVs to support assay data of live VSV-S-M^pro^ dose-responses

We sought to perform additional dose-response experiments using an orthogonal assay employing VSV particles pseudotyped with SARS-CoV-2 spike protein (XBB.1.5 variant). We cloned two spike variants from the pool generated during virus generation and selection experiments with RQ-01. The spike gene of the codon-optimized full-length spike XBB.1.5 was amplified via PCR with the following primers (“T807_SXBB1.5_CO_for“ and „T807_SXBB1.5_CO_rev“ from **Table S1**) from an expression plasmid encoding this gene (SAVE Consortium). We introduced amino acids substitutions through mutagenic Gibson assembly using the primers listed in **Table S1**.

After cloning the spike variant plasmids, we transfected 293T cells with these plasmids via calcium phosphate transfection (as described in “**viral rescue**”), where instead of using chloroquine, we used DMEM without FCS supplementation. One day prior transfection, we seeded two 10 cm dishes at a confluence of 2×10^6^ cells/dish for each spike variant. After addition of the master mix to the cells, we incubated the dishes for 8 hours at 37°C. Then, we removed the medium without FCS with a vacuum pump and replaced it with fully supplemented DMEM. Two days after transfection, we infected all dishes with MOI = 3 with a VSV variant where the VSV-G gene was replaced by GFP. These single-round infectious VSV particles are pseudotyped with the glycoprotein of the lymphocytic choriomeningitis virus (LCMV-GP) and can infect 293T cells. After 90 minutes, we removed the medium containing VSV-ΔG again with a vacuum pump. We carefully washed 293T cells once with PBS and then added fresh, fully supplemented DMEM containing rabbit serum neutralizing the remaining VSV-ΔG-GFP (pseudotyped with the GP from LCMV) in the supernatant and incubated for 24 to 30 hours until GFP signal was abundant under the fluorescent microscope. We also transfected an expression plasmid encoding VSV-G and pseudotyped VSV with G to use it as negative control in the dose-response assays. If GFP signal was present, we harvested the supernatant, filtered through a 45 μm filter and aliquoted in 1.5 mL microcentrifuge tubes.

We then used these spike-pseudotyped VSVs to perform dose-response experiments with RQ-01 to assess the loss of susceptibility against spike HR1/HR2 variants. In brief, we seeded 1.8×10^4^ 293T A4 (↑ACE2) cells in 96 well plates one day before infection. At the day of infection, we prepared pseudotyped virus dilutions to achieve approximately 200 GFP spots/well in the untreated condition. We serially diluted RQ-01 in the virus-containing medium in a half-log dilution fashion reaching seven concentration conditions and leaving the 8^th^ untreated. Concentration ranges either started from 1 or 0.5 μM. We read out the GFP signal after 16 hours post infection.

We retrieved spot count data by using the FluoroSpot X Suite on our ELISpot reader. We then normalized the spot count from 0 to 1, determined IC_50_ values using the same built-in equation from GraphPad Prism used before (“Sigmoidal, 4PL, X is concentration”) and finally calculated IC_50_ fold-changes.

### Structure collection for pre- and post-fusion stability investigation

To evaluate the stability effect of mutations on both the pre- and post-fusion conformation of the SARS CoV-2 omicron spike protein, structural information is necessary. Experimentally determined structure models for both states were taken from PDB ID 7TEI [68] for the pre-fusion conformation of HR1 domain, PDB ID 2FXP [48] for the pre-fusion conformation of HR2 domain and 7TIK [44] for the post-fusion conformation.

### Computational modeling and evaluation of the mutations

The structure models were relaxed using FastRelax [69] as implemented in the Rosetta molecular modeling suite [70,71]. Backbone flexibility was achieved via the BackrubProtocol, [72] which samples conformational changes. This approach enabled the exploration of alternative low-energy backbone conformations while preserving the overall fold of the protein [73]. Each Backrub run consisted of 10000 independent backrub trial moves with a Monte Carlo criterion at a temperature factor of 0.3. The protocol was executed in 1000 independent iterations, each starting from the relaxed structure, to generate protein structures representing plausible backbone variability to account for backbone flexibility.

### Calculation of thermostability profiles

Thermostability profiles of the generated protein structures were calculated using THERMOMPNN [74], a message passing neural network based on the PROTEINMPNN [75] architecture combined with a transfer learning block, which was trained on the Megascale stability dataset [76] to predict stability changes upon amino acid changes. We conducted a two-state THERMOMPNN calculation following a previously established protocol (Riccabona et al., in preparation). This protocol requires both the pre- and the post fusion conformation of the SARS CoV-2 omicron spike protein. With this protocol, relative stability challenges in between pre- and post-fusion conformation can be evaluated. As we wanted to evaluate, which mutations stabilize the post-fusion conformation compared to the pre-fusion, we used the following formula of the relative thermostability profile:

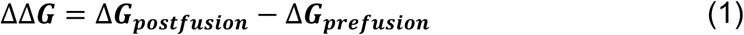

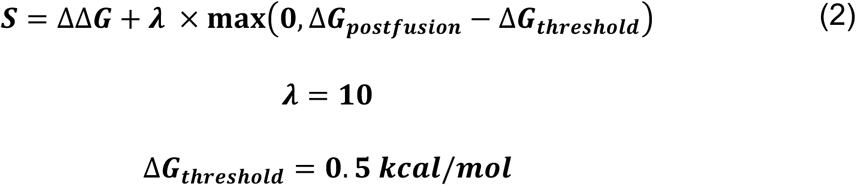

***λ*** being a weighting factor and Δ***G***_***threshold***_ representing a destabilizing tolerance threshold. This formula penalized mutations that increased the free energy of the post-fusion conformation beyond the defined threshold.

### Hardware overview (TTMD simulations)

Molecular modeling tasks, such as complex preparation, in silico mutations, system setup for molecular dynamics (MD), and simulation analysis, were carried out on a Linux workstation running Ubuntu 22.04 LTS and shipping a 24-core Intel Core i9-14900K 6.0 GHz processor. Molecular dynamics simulations were performed on an in-house Linux GPU cluster comprising 30 NVIDIA GPUs ranging from GTX1080Ti to RTX4090.

### Structure Preparation

Structure preparation and computational workflows were conducted using the Molecular Operating Environment (MOE) 2024 software suite [77]. The three-dimensional (3D) structure of the SARS-CoV-2 six-helix bundle (6-HB) was retrieved from the PDB [78] (PDB ID 7TIK). To mimic the RQ-01 peptide, the D chain of the complex was truncated on the C-terminal residue D1168, and residues K1181 and S1188 were modified and cross-linked with an 11-carbon tether staple. Relevant amino acid changes were added using the “Protein Builder” tool in MOE, followed by energy minimization with the Amber 14:EHT force field. The “Structure Preparation” module was employed during the preparation process, fixing eventual inconsistencies. N/C-termini were capped with acetyl and N-methylamide, respectively, and hydrogens were added through “Protonate 3D” following the most probable tautomeric and protomeric state at PH 7.4.

### System setup for MD simulations

All the complexes obtained were further prepared for MD simulation exploiting the Amber Tools 22 suite [79]. Parameters were attributed according to the ff14SB force field [80]. Each complex was solvated in a rectangular base box, keeping at least 15 Å of padding between the box edges, using the TIP3P water model [81]. Sodium and chloride ions were added to neutralize the net charge of the system, reaching a concentration of 0.154 M. To eliminate steric clashes and unfavorable contacts, an initial energy minimization of 500 steps was carried out using the conjugate gradient algorithm, before the equilibration phase.

Before running the production simulations in the canonical NVT ensemble, a two-step equilibration protocol was performed. The first step involved a short simulation (0.1 ns) in the NVT ensemble, during which harmonic positional restraints of 5 kcal·mol⁻¹·Å⁻² were applied to all the atoms of the complex, allowing solvent and ions to move freely. This was followed by an isothermal–isobaric ensemble (NPT) simulation of longer duration (0.5 ns), where restraints were maintained only on the backbone atoms. Throughout both stages, the temperature was held constant at the target value using a Langevin thermostat [82], while, during the NPT phase, the pressure was regulated at 1 atm through a Monte Carlo barostat [83].

All molecular dynamics simulations were carried out using the ACEMD 3.8.2 engine [84], which is based on the open-source OpenMM library [85]. A 2 fs integration timestep was employed, and the M-SHAKE algorithm [86] was used to constrain bonds involving hydrogen atoms. Electrostatic interactions were computed using the particle-mesh Ewald (PME) method [87] with cubic spline interpolation, while a 9.0 Å cutoff was applied to Lennard–Jones interactions.

### Thermal titration molecular dynamics (TTMD) simulations

Thermal Titration Molecular Dynamics (TTMD) [88] is an enhanced sampling protocol originally introduced for the qualitative assessment of protein–ligand complex stability, which has already been applied to study the alteration of complex stability caused by mutations [20].

This method consists of a sequence of short molecular dynamics simulations defined as “TTMD-steps” performed at progressively increasing temperatures, thereby accelerating unbinding events compared to conventional MD. In this study, the ramp ranged from 300 K to 450 K, and each step of 10 ns was performed with a temperature increase of 10 K.

To evaluate the persistence of the native binding mode during the simulation, an interaction fingerprint–based scoring function (IFPcs) [89] was employed. For each frame, the interaction between the RQ-01 model and the 5-helix bundle (5-HB) was encoded into an interaction fingerprint using the ProLIF Python package. [90] Each frame-fingerprint was compared to the reference fingerprint obtained from the last frame of equilibration through the cosine similarity. The resulting similarity was multiplied by −1, yielding scores ranging between −1, indicating the full conservation of the binding mode, and 0 when the native contacts are completely lost. At the end of each TTMD-step, the average score was calculated, and if this value exceeded −0.05, the simulation was terminated early, indicating disruption of the native binding mode.

The overall stability of each complex was further quantified through the MS coefficient, defined as the slope of the line connecting the origin of the titration profile graph and its final point, which is the average IFPcs value for the last step of the simulation.

Values close to 0 indicate conservation of the binding mode, whereas values near 1 indicate a loss of native interactions. For each complex, five independent TTMD runs were performed, and the MS score was averaged across three replicates, discarding the highest and lowest values. In addition, a titration timeline plot was generated, reporting the evolution of IFPcs, thus providing a comprehensive view of binding stability during the simulation under thermal stress.

## Acknowledgements

We gratefully acknowledge all data contributors, i.e., the authors and their originating laboratories responsible for obtaining the specimens, and their submitting laboratories for generating the genetic sequence and metadata and sharing via the GISAID Initiative, on which this research is based. We want to acknowledge the company Red Queen Therapeutics (RQT), which has provided the peptide fusion inhibitor RQ-01. We want to thank the SAVE consortium and Janine Kimpel for providing a codon-optimized XBB.1.5 spike expression plasmid. We are grateful to Janine Kimpel for providing the rabbit serum used to produce the VSV-Spike pseudotypes. We are thankful to Dr. med. David Bante for providing the SARS-CoV-2 XBB.1.5 cDNA used to generate our dual-chimeric VSV and developing the scripts used to analyze the ONT data. The MMS lab is grateful to Chemical Computing Group, OpenEye, and Acellera for the scientific and technical partnership. MMS lab thankfully acknowledges the support of NVIDIA Corporation with the donation of the Titan V GPU used for this research.

## Author contributions

**E.H.:** conception and supervision of the study, funding acquisition, generation of the dual-chimeric virus, data acquisition and analysis, original manuscript drafting; **F.C.:** funding acquisition, performing experiments (dose-response assays with live and pseudotyped viruses and serial virus passages), ONT sequencing, data acquisition and analysis, figure preparation, original manuscript drafting; **A.D.:** performing classical molecular dynamics and TTMD analyses, figure preparation, manuscript drafting, and proofreading; **A.F.:** production of large-scale plasmid DNA, generation of pseudotyped viruses, and performing pseudotyped virus assays; **S.R.:** performing experiments, carrying out upstream tasks prior to ONT sequencing (viral RNA isolation, cDNA, PCR, etc…), assistance with virological data acquisition and analysis; **J.R.R.:** ThermoMPNN analyses, manuscript drafting, proofreading; **A.B.:** carrying out upstream tasks prior to ONT sequencing, assistance with serial virus passages, and proofreading; **L.K.:** provision of anti-XBB.1.5 serum for neutralization assays; **L.S.:** data interpretation throughout the project and manuscript proofreading; **C.T.S.:** supervision of ThermoMPNN analyses, contribution to data interpretation. **S.M.:** supervision of molecular dynamics and TTMD analyses, contribution to data interpretation. **G.G.:** supervision of the study, contribution to data interpretation, assistance with manuscript drafting and proofreading.

## Funding

This work was funded by the Austrian Science Fund (FWF) grant P35148 with DOI 10.55776/P35148 and by the State of Tyrol in the Tyrolean Young Researchers Funding program (Tiroler Nachwuchsforscher*innenfoerderung). Some of the research material required for this work, such as Nanopore sequencing flow cells and reagents, was funded by RQT. The salary of E.H. was covered by the Global Fellowship Program of the King Abdullah University of Science and Technology (KAUST). J.R.R. and C.T.S were supported by the Federal Ministry of Education and Research of Germany and by the Sächsiches Staatsministerium für Wissenschaft Kultur und Tourismus in the program Center of Excellence for AI-research “Center for Scalable Data Analytics and Artificial Intelligence Dresden/Leipzig”, project identification number ScaDS.AI. For open access purposes, the author has applied a CC BY public copyright license to any author accepted manuscript version arising from this submission.

## Competing interests

The data in this work was generated in the context of a drug approval process with the US Federal Drug Administration (FDA) for the clinical licensing of the compound RQ-01.

## Data availability

TTMD and MD data can be found on Zenodo: https://doi.org/10.5281/zenodo.18116971. TTMD code is released under a permissive MIT license and available free of charge at github.com/molecularmodelingsection/TTMD.

## Supplemental information

**Supplemental Figure 1.**
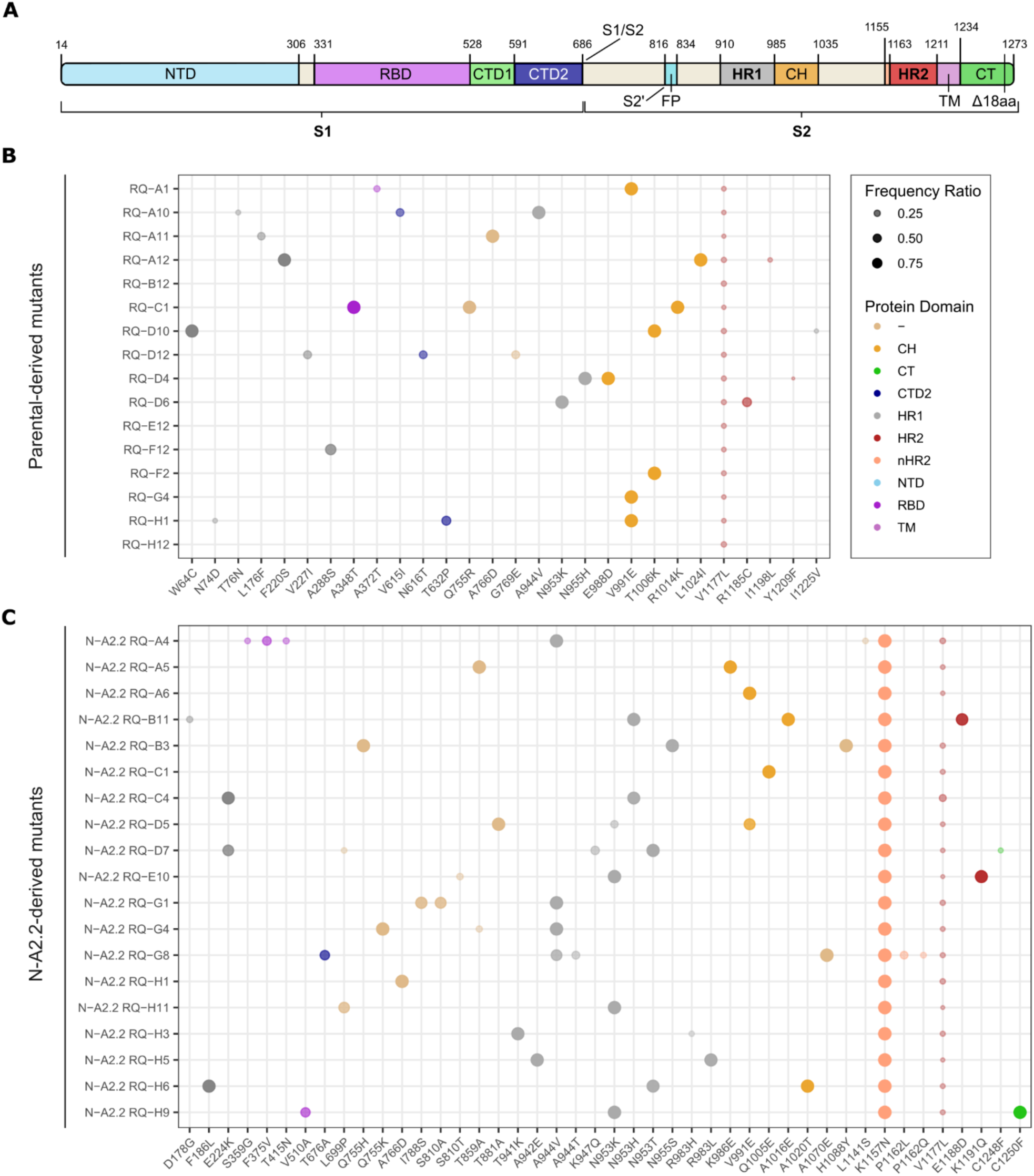
ONT sequencing of VSV-S-M^pro^-derived viruses after ten passages with RQ-01 revealed AA substitutions in the spike gene. Plots in this figure provide the following information: sample name, AA substitution(s) of each sample and the population fraction/frequency ratio of the sequence carrying the single nucleotide variation (SNV) that leads to that specific AA substitution (from 0 to 1). The AA substitutions are plotted on the x-axis and the sample names are plotted on the y-axis. *Sample nomenclature*: indication of the VSV-S-M^pro^ variant used to generate mutants (parental or N-A2.2); the inhibitor used to generate spike variants (here only RQ-01) and the well from which the sample was collected after 10 passages (A10, D6, etc.). Example name of a variant generated from VSV-S-M^pro^ is “RQ-A10”, while the name of a variant generated from VSV-S-M^pro^-E166V (N-A2.2) is “N-A2.2 RQ-H3”. Color-coding in the dot-plot is assigned according to the spike protein domains shown in (**A**). **A)** Schematic view of the spike (S) protein and domains [43]. “CH” = central helix; “CT” = cytoplasmic tail; “CTD2” = carboxy-terminal domain 2; “FP” = fusion peptide; “HR1” = heptad-repeat 1; “HR2” = heptad repeat 2; “NTD” = amino-terminal domain; “RBD” = receptor binding domain; “TM” = transmembrane anchor. Specific naming used in in this figure: “nHR2” = near HR2 and “-” = no assigned domain. **B)** ONT sequencing results of the samples collected from serial passaging experiments of parental VSV-S-M^pro^ with RQ-01. **C)** ONT sequencing results of the samples collected from serial passaging experiments of N-A2.2 (VSV-S-M^pro^-E166V) with RQ-01. The legend box indicates the frequency ratio (metric already shown in **Figure 2** and **3**). Small and faded circles indicate a low population/frequency ratio of specific mutants in the sequences pool from a specific sample, whereas big and vivid circles indicate the opposite. For both (**B**) and (**C**) we did not include R682W in the plot as it was present in all the samples before passaging.

**Supplemental Figure 2.**
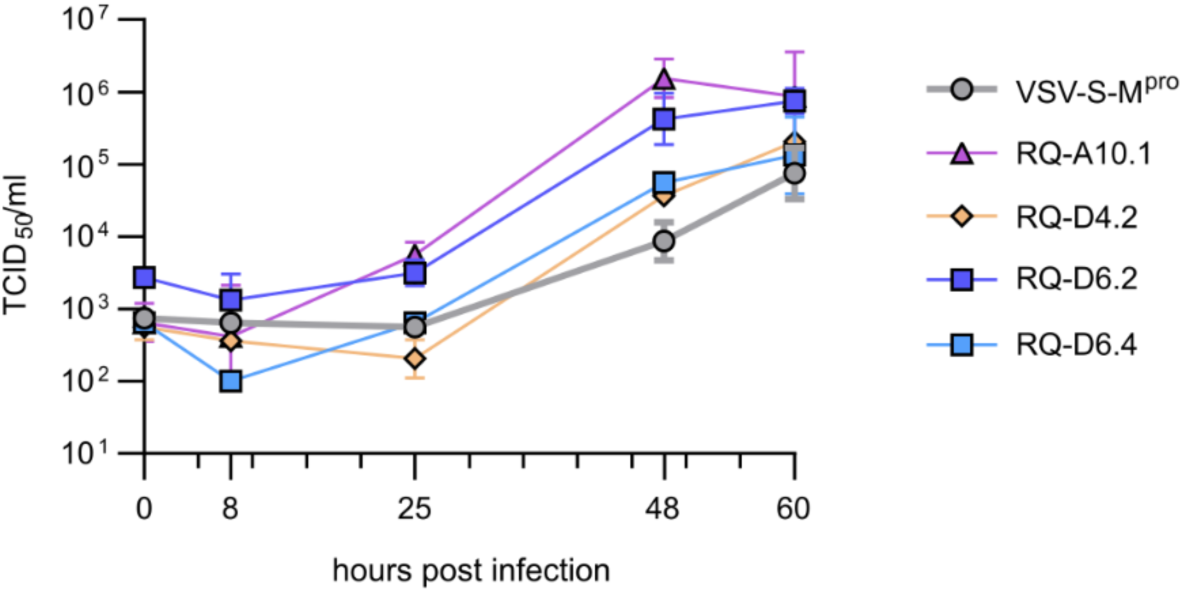
Conventional multistep viral growth kinetic supports the results obtained with mWasabi kinetic experiments. Multi-step growth kinetics experiment with the variants VSV-S-M^pro^ parental virus, RQ-A10.1, RQ-D4.2, RQ-D6.2 and RQ-D6.4. Titers at different time points are represented as TCID_50_/mL. Shorter ticks represent five hours intervals. Data are presented as geometric mean ± SD of 2 technical replicates per condition.

**Supplemental Figure 3.**
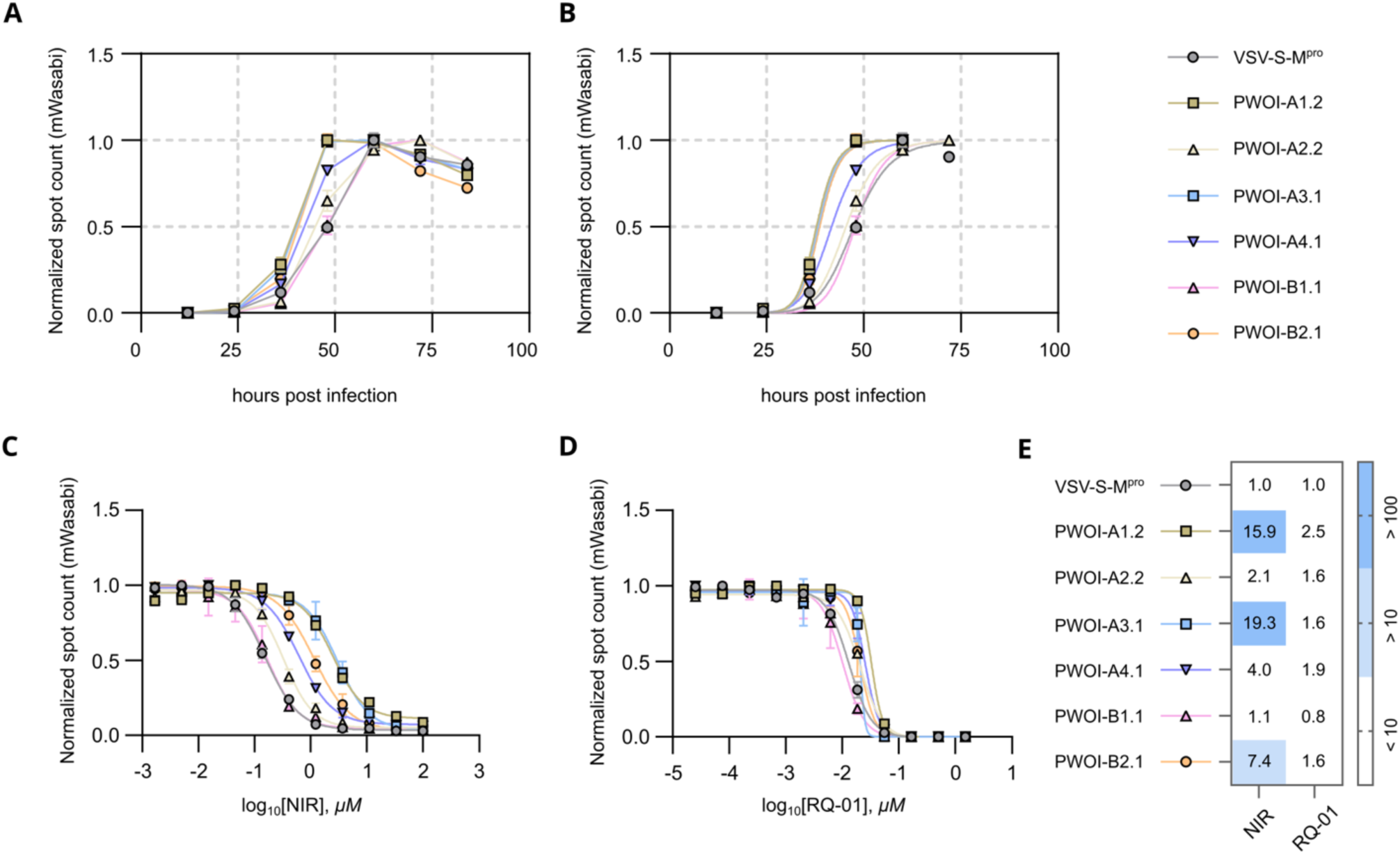
Viruses passaged without inhibitor (‘PWOI’) acquire cell culture adaptations. **A)** mWasabi signal growth over time. Data are presented as mean ± SD of 4 technical replicates per condition. **B)** Non-linear regression of mWasabi signal growth over time (**panel A**). Data are presented as mean ± SD of 4 technical replicates per condition. **C,D)** Dose-response experiments of PWOI variants against the inhibitors NIR (**C**) and RQ-01 (**D**). mWasabi spot count was measured at 60 hours post infection. Data is presented as mean ± SEM of 4 technical replicates per condition. The curve representing the variant PWOI-A3.1 in the RQ-01 dataset is presented as mean ± SEM of 3 technical replicates per condition. **E)** Heat map including the calculated IC_50_ fold-changes of all the tested VSV-S-M^pro^ spike variants compared to the original virus (as internal control), for both inhibitors. Symbols and lines of the dose response curves are placed to the side of the VSV-S-M^pro^ variant names.

**Supplemental Figure 4.**
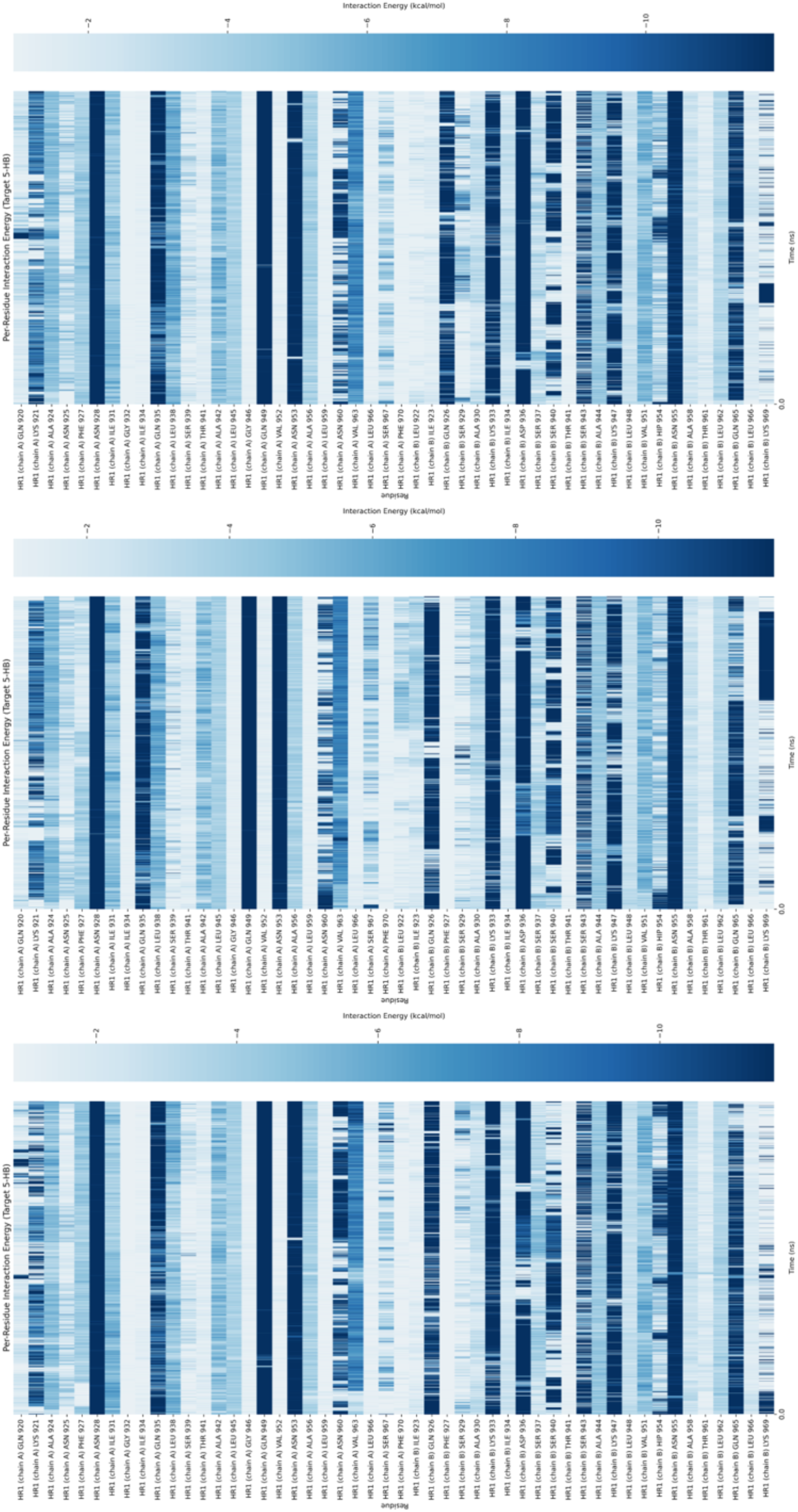
Per-residue interaction energy plots of the three MD simulations of the stapled complex (stap-HR1/HR2).

**Supplemental Figure 5.**
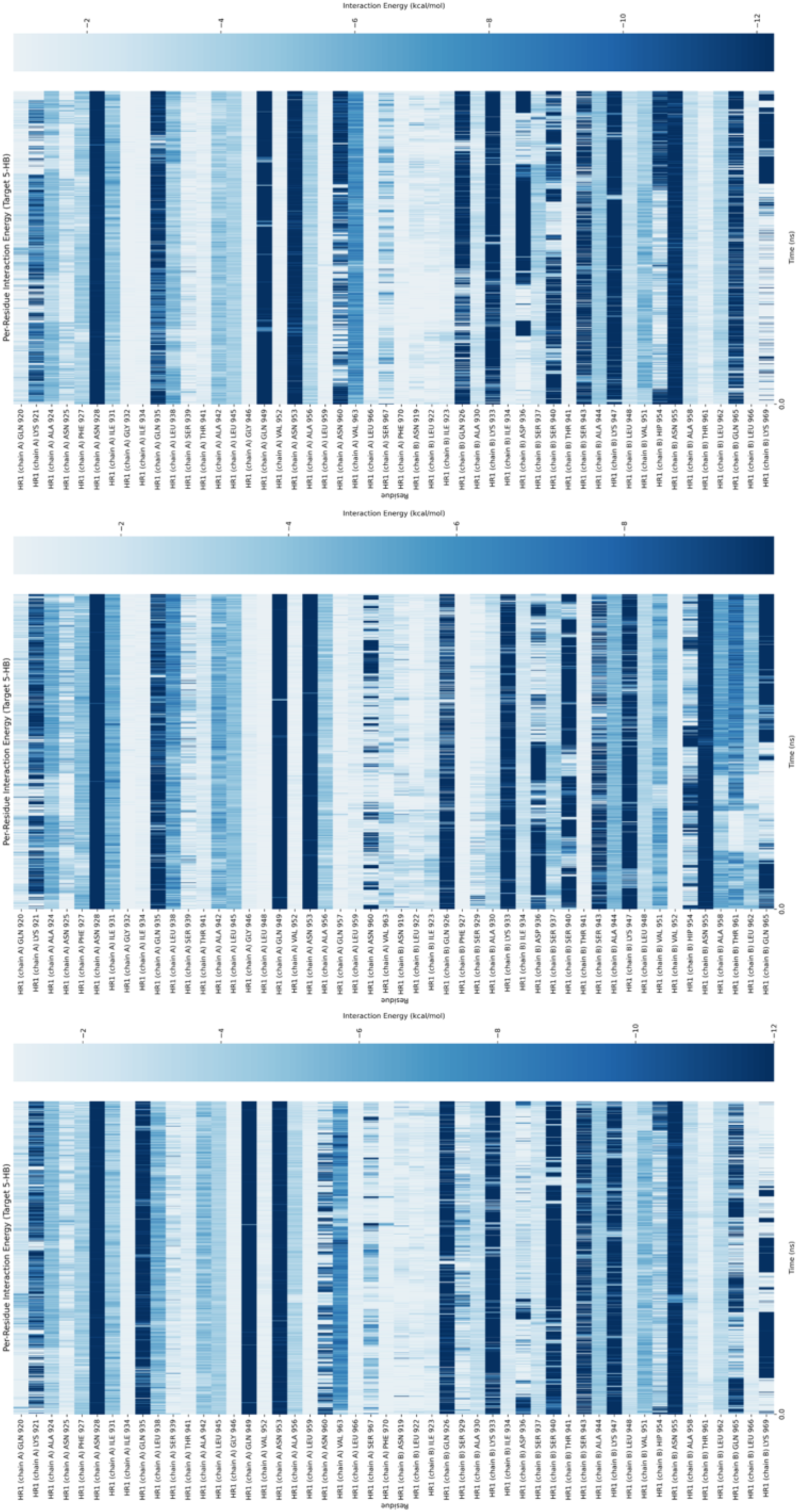
Per-residue interaction energy plots of the three MD simulations of the non-stapled complex (nstap-HR1/HR2).

**Supplemental Figure 6.**
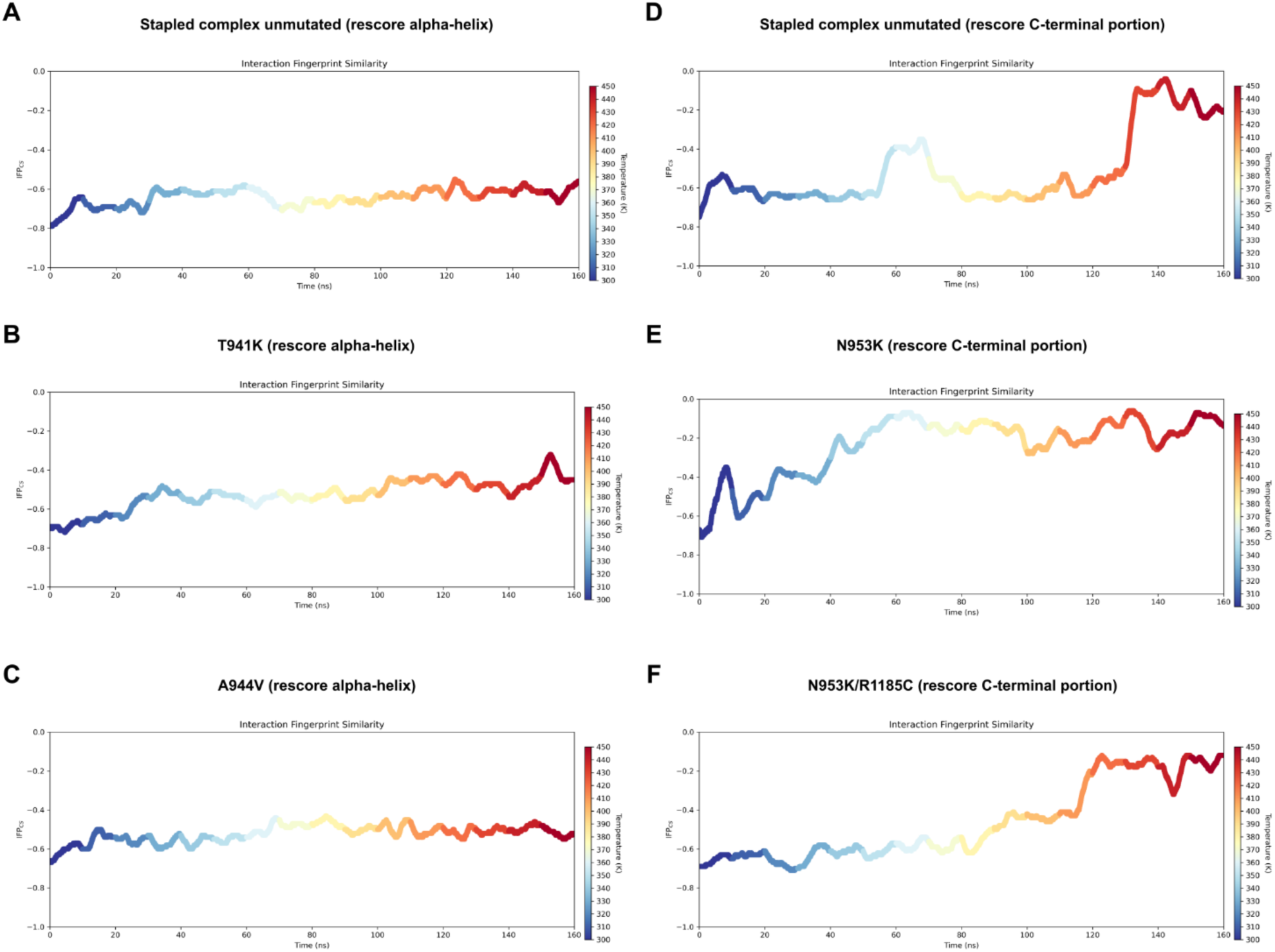
**A)** Titration timeline plots of the stapled complex unmutated and the variants T941K (**B**) and A944V (**C**) after rescoring of the alpha-helix. The MS rescoring for A944V and T941K considered only the alpha helical portion of HR2 (I1179-S1196), since it is directly in contact with these residues. This rescoring amplified the MS score difference between these stap-HR1/HR2 mutants and the unmutated stap-HR1/HR2, emphasizing their higher predicted instability (**Supplemental Table S4**). **D)** Titration timeline plots of the stapled complex unmutated and the variants N953K (**E**) and N953K/R1185C (**F**) after rescoring of the C-terminal portion. The MS scores of both variants indicate a slightly higher predicted complex instability for the mutants compared to the unmutated complex, with mean values of 0.00550 for N953K and 0.00615 for N953K/R1185C, versus 0.0053 for the unmutated stap-HR2 (**Supplemental Table S5**).

**Supplemental Figure 7.**
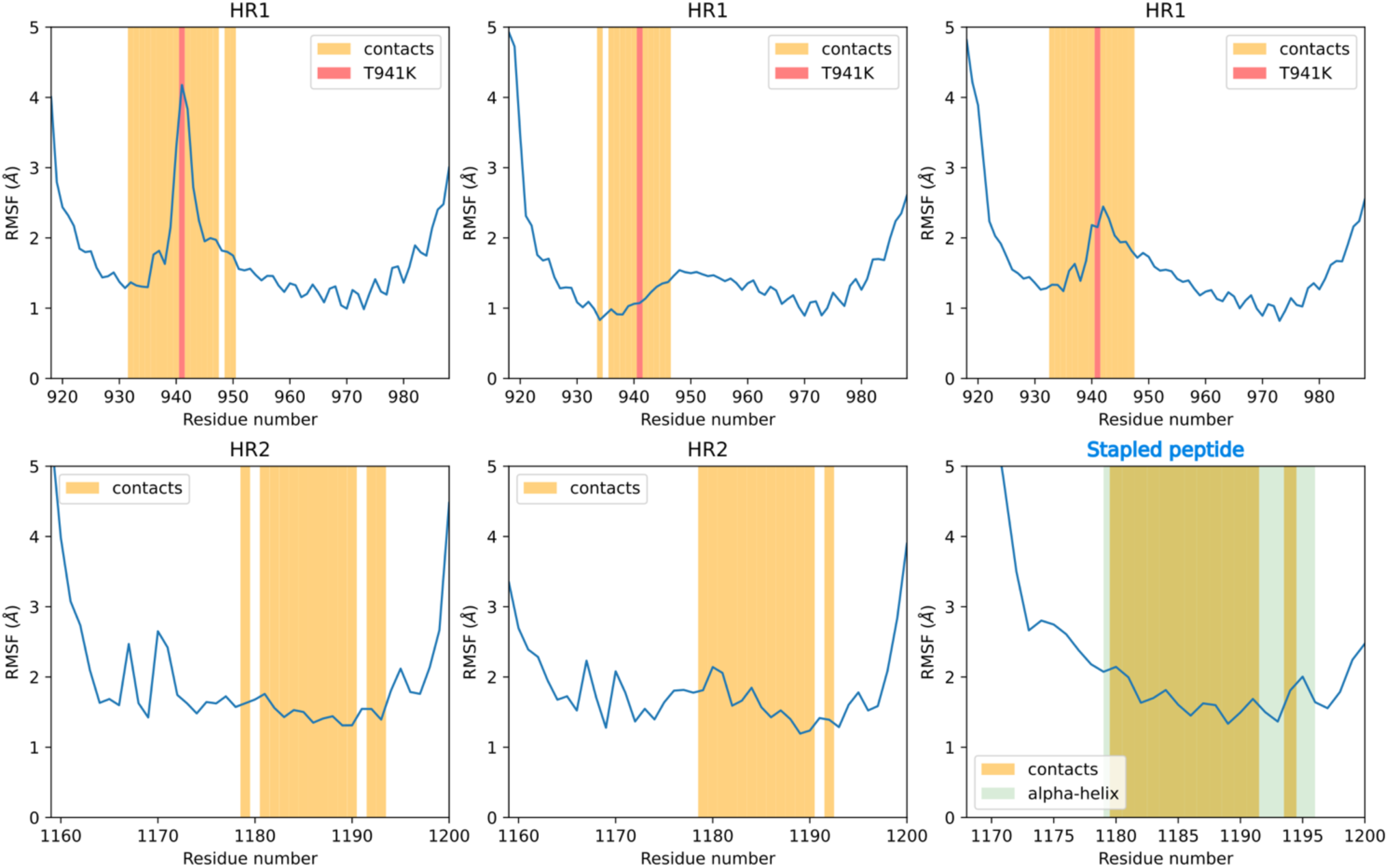
RMSF profile of T941K TTMD representative simulation after rescoring. RMSF profile of T941K. T941K is indicated by a red line, its contacts in yellow, and the alpha-helix portion of RQ-01 in light lime. From left to right: top, chain A - chain B - chain C of PDB 7tik; bottom, chain E - chain F - chain D (stapled HR2 decoy, RQ-01). In this simulation, K941 displayed a highly dynamic behavior primarily driven by electrostatic interactions. This substitution allows engagement with multiple residues and causes a distortion of the local alpha-helix geometry that leads to a disruption of native interactions. We observed that by interacting with D936, K941 breaks the R1185-D936 salt bridge and occasionally competes with R1185 for E1182, thereby perturbing the E1182-K947 salt bridge as well. We observed also occasional interactions with I1179 and Q1180 that altered hydrogen bonds involving S940 and S943.

**Supplemental Table 1.**
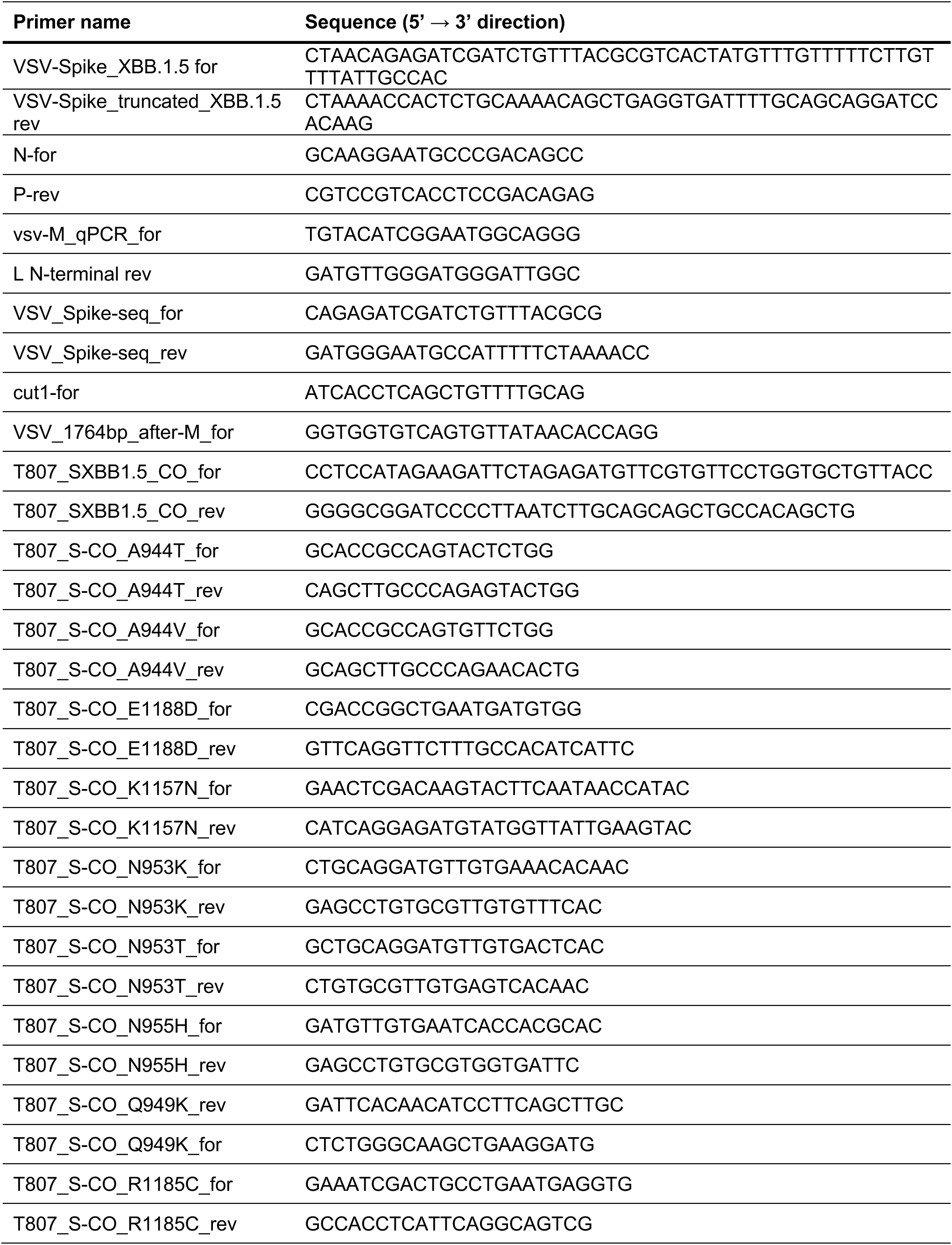
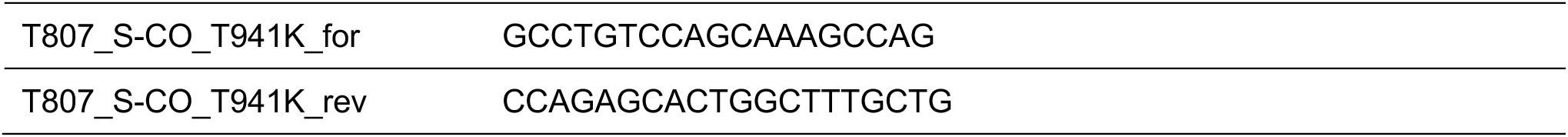
Primers used for VSV-PmWasabi-Spike-M^pro^ cloning, ONT sequencing and expression plasmids for VSV pseudotypes.

**Supplemental Table 2.** GISAID prevalence of spike substitutions detected after ONT sequencing (as of 18^th^ November 2025). The search was done selecting “complete” and “low coverage excluded” in EpiCoV [45–47]. “All-variants” refers to the number of entries across all SARS-CoV-2 variants. “Only XBB.1.5” refer to the number of entries of XBB.1.5 variant only. “Ratio all-variants/XBB.1.5” refers to the ratio between entries across all variants and entries from XBB.1.5 only.

| Substitution | All-variants | Only XBB.1.5 | Ratio all-variants/XBB.1.5 | Domain |
| --- | --- | --- | --- | --- |
| A4T | 0 | 0 | - | NTD |
| L54S | 184 | 5 | 35.80 | NTD |
| F58L | 228 | 26 | 7.77 | NTD |
| W64C | 1728 | 46 | 36.57 | NTD |
| N74D | 1662 | 32 | 50.94 | NTD |
| T76N | 863 | 75 | 10.51 | NTD |
| S151N | 595 | 54 | 10.02 | NTD |
| L176F | 29942 | 909 | 31.94 | NTD |
| D178G | 1267 | 7 | 180.00 | NTD |
| F186L | 36302 | 381 | 94.28 | NTD |
| F220S | 43 | 2 | 20.50 | NTD |
| E224K | 86 | 0 | - | NTD |
| V227I | 1406 | 57 | 23.67 | NTD |
| A288S | 864 | 105 | 7.23 | NTD |
| A348T | 1671 | 1149 | 0.45 | RBD |
| S359G | 296 | 15 | 18.73 | RBD |
| A372T | 173 | 2 | 85.50 | RBD |
| F375V | 0 | 0 | - | RBD |
| K386R | 30 | 0 | - | RBD |
| K386R | 30 | 0 | - | RBD |
| T415N | 666 | 26 | 24.62 | RBD |
| N417Y | 0 | 0 | - | RBD |
| V510A | 10 | 2 | - | RBD |
| D571G | 99 | 9 | 4.00 | CTD1 |
| V615I | 1755 | 37 | 10.00 | CTD2 |
| N616T | 7 | 0 | 46.43 | CTD2 |
| V622D | 330 | 5 | - | CTD2 |
| T632P | 7 | 1 | 65.00 | CTD2 |
| G648S | 3 | 0 | 6.00 | CTD2 |
| C649W | 1 | 0 | - | CTD2 |
| T676A | 559 | 6 | - | CTD2 |
| R682W | 420 | 3 | 92.17 | S1/S2 |
| R683Q | 2908 | 15 | 139.00 | S1/S2 |
| S689N | 687 | 7 | 192.87 | - |
| L699P | 15 | 1 | 97.14 | - |
| Q755H | 178 | 81 | 14.00 | - |
| Q755K | 8 | 0 | 1.20 | - |
| Q755R | 34 | 1 | - | - |
| A766D | 15 | 0 | 33.00 | - |
| G769E | 197 | 8 | - | - |
| I788S | 1 | 0 | 23.63 | - |
| P793L | 912 | 15 | - | - |
| S810A | 382 | 6 | 59.80 | - |
| S810T | 61 | 1 | 62.67 | - |
| N824D | 314 | 3 | 60.00 | FP |
| T859A | 385 | 40 | 103.67 | - |
| T881A | 385 | 1 | 8.63 | - |
| A924S | 1944 | 8 | 384.00 | HR1 |
| T941K | 15 | 1 | 242.00 | HR1 |
| A942E | 11 | 1 | 14.00 | HR1 |
| A944T | 23 | 0 | 10.00 | HR1 |
| A944V | 43 | 1 | - | HR1 |
| K947Q | 6 | 0 | - | HR1 |
| Q949K | 39 | 1 | 38.00 | HR1 |
| N953H | 0 | 0 | - | HR1 |
| N953K | 9 | 0 | - | HR1 |
| N953T | 4 | 0 | - | HR1 |
| N955H | 44 | 0 | - | HR1 |
| N955S | 3 | 0 | - | HR1 |
| R983H | 4 | 0 | - | HR1 |
| R983L | 534 | 0 | - | HR1 |
| K986E | 19 | 1 | 18.00 | CH |
| E988D | 52 | 0 | - | CH |
| V991E | 2 | 0 | - | CH |
| Q1005E | 1 | 0 | - | CH |
| T1006K | 1 | 0 | - | CH |
| R1014K | 1255 | 42 | 28.88 | CH |
| A1016E | 5 | 0 | - | CH |
| A1020T | 562 | 12 | 45.83 | CH |
| L1024I | 155 | 13 | 10.92 | CH |
| A1070E | 107 | 0 | - | - |
| H1088Y | 769 | 23 | 32.43 | - |
| L1141S | 71 | 0 | - | - |
| K1157N | 1134 | 6 | 188.00 | nHR2 |
| P1162L | 27453 | 474 | 56.92 | nHR2 |
| P1162Q | 359 | 32 | 10.22 | nHR2 |
| V1177L | 1618 | 17 | 94.18 | HR2 |
| R1185C | 467 | 5 | 92.40 | HR2 |
| E1188D | 1702 | 35 | 47.63 | HR2 |
| K1191Q | 295 | 2 | 146.50 | HR2 |
| I1198L | 33 | 1 | 32.00 | HR2 |
| Y1209F | 49 | 0 | - | HR2 |
| I1225V | 1611 | 46 | 34.02 | TM |
| C1248F | 3806 | 32 | 117.94 | CT |
| C1250F | 5340 | 41 | 129.24 | CT |

**Supplemental Table 3.**
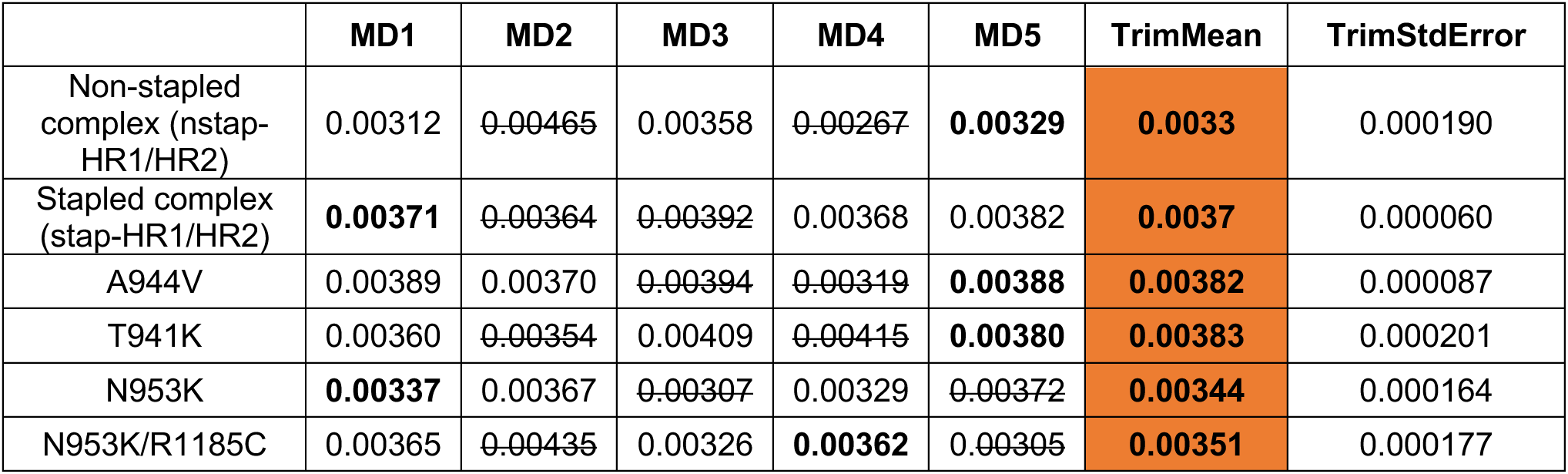
Overall TTMD results. In bold is indicated the representative simulation and the mean MS score highlighted in orange.

**Supplemental Table 4.**
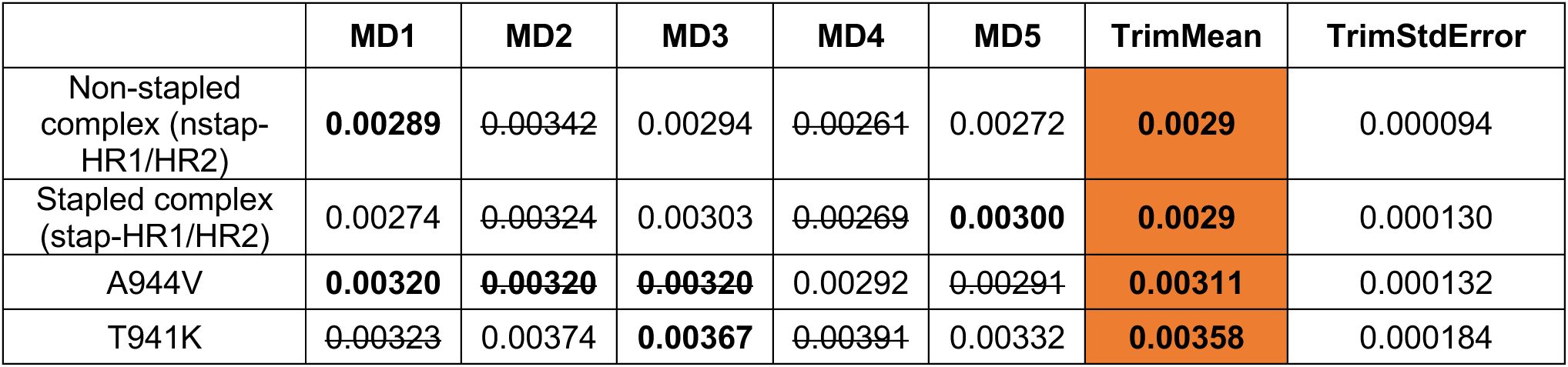
TTMD rescoring results on the alpha-helix portion of RQ-01 (I1179-S1196). In bold is indicated the representative simulation and the mean MS score highlighted in orange.

**Supplemental Table 5.**
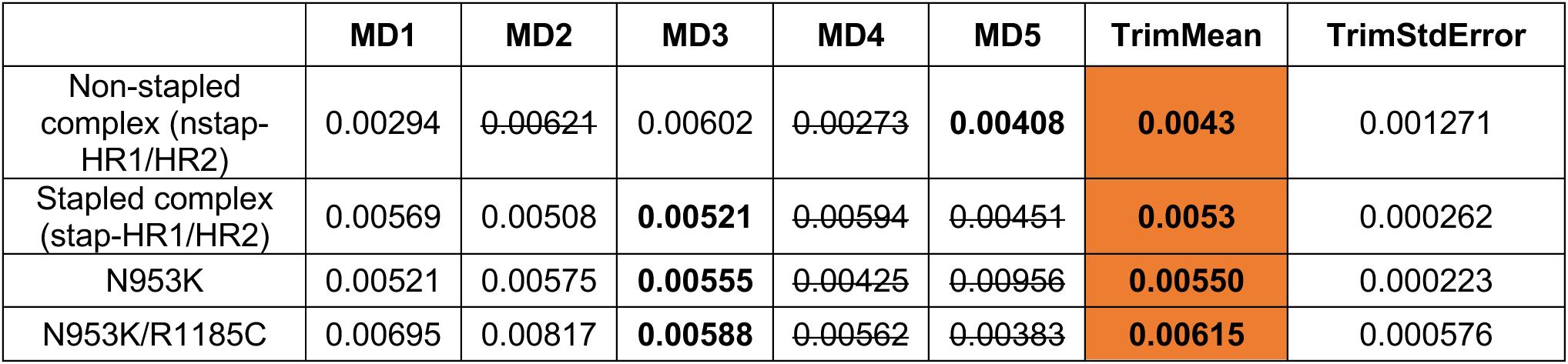
TTMD rescoring results on the C-terminal portion of RQ-01 (D1168-N1178). In bold is indicated the representative simulation and the mean MS score highlighted in orange.

## Legend for Supplemental Videos 1-4

**Supplemental Video 1. TTMD simulation video of the T941K variant. Additional comment:** a closer inspection of the simulation reveals that the T941K mutant is highly unstable.

**Supplemental Video 2. TTMD simulation video of the A944V variant. Additional comment:** we carried out a rescoring for A944V (**Supplemental Figure S6C**), obtaining a mean MS score of 0.00311 (vs 0.0029 of the unmutated stap-HR1/HR2), indicating increased predicted instability. The representative simulation had an MS score of 0.00320 (vs 0.00300 of the unmutated stap-HR1/HR2). The substitution from A944 to V944 affects the interaction patterns of nearby residues due to the higher steric hindrance of valine. The most affected residue is E1182, which interacts with R1185 as a result of this substitution. Since now R1185 is engaged with E1182, the alpha helix portion of RQ-01 can drift away from the substituted HR1, causing a disruption of key the salt bridges like R1185-D936 and E1182-K947 and hydrogen bonds of S943 and S940 with E1182.

**Supplemental Videos 3-4. TTMD simulation video of the N953K and N953K/R1185C variants, respectively. Additional comment:** these simulations highlight how the shared N953K mutation locally alters the stap-HR1/HR2 binding mode. Here, N953 forms a hydrogen bond with the backbone of V1177, anchoring HR1 and HR2, ensuring optimal orientation towards hydrophobic residues. Upon substitution to K953, the N953-V1177 hydrogen bond is lost and the local flexibility of RQ-01 is enhanced in the A1174-N1178 stretch. The resulting RMSD values were 3.4056 Å for N953K, 3.03909 Å for N953K/R1185C, and 2.2219 Å for the stap-HR1/HR2. We observed a similar trend when extending the analysis to the entire C-terminal tail (D1168-N1178), obtaining higher results for N953K (5.59 Å) and N953K/R1185C (4.61 Å) compared to the stap-HR1/HR2 (4.02 Å). We observed that while the local impact of the N953K mutation on the (RQ-01)-(5-HB) interface is clear, the R1185C mutation, due to the high distance from RQ-01, does not appear to have a direct effect on the stability of the complex under the simulated temperature ramp.

## Notes

https://doi.org/10.5281/zenodo.18116971

https://github.com/molecularmodelingsection/TTMD

